# The natural adaptation and human selection history of African sheep genomes

**DOI:** 10.1101/2021.11.17.469022

**Authors:** Abulgasim M. Ahbara, Christelle Robert, Adebabay Kebede, Ayelle Abeba, Suliman Latairish, Mukhtar Omar Agoub, Hassan H. Musa, Pam Wiener, Emily Clark, Olivier Hanotte, Joram M. Mwacharo

**Affiliations:** Department of Zoology, Faculty of Sciences, Misurata University, Misurata, Libya; School of Life Sciences, University of Nottingham, University Park, Nottingham, UK; Small Ruminant Genomics, International Center for Agricultural Research in the Dry Areas (ICARDA), Addis Ababa, Ethiopia; LiveGene, International Livestock Research Institute (ILRI), Addis Ababa, Ethiopia; Centre for Tropical Livestock Genetics and Health (CTLGH), The Roslin Institute, University of Edinburgh, UK; LiveGene—CTLGH, International Livestock Research Institute (ILRI) Ethiopia, Addis Ababa, Ethiopia; Amhara Regional Agricultural Research Institute, Bahir Dar, Ethiopia; Debre Berhan Research Centre, Debre Berhan, Ethiopia; Department of Animal Production, Faculty of Agriculture, Misurata University, Misurata, Libya; Agricultural Research Center, Misurata, Libya; Faculty of Medical Laboratory Sciences, University of Khartoum, Sudan; Animal and Veterinary Sciences, SRUC, The Roslin Institute Building, Midlothian Edinburgh, UK

**Keywords:** Diversity, Demographic history, Fat-rumped, Fat-tailed, Ovis aries, Thin-tailed

## Abstract

African sheep manifest diverse but distinct physio-anatomical traits which are the outcomes of natural- and human-driven selection. Here, we generated 34.8 million variants from 150 indigenous African sheep genomes sequenced at an average depth of ∼54x for 130 samples (Ethiopia, Libya) and ∼10x for 20 samples (Sudan), representing sheep from diverse environments, tail morphology and post-Neolithic introductions to Africa. Phylogenetic and model-based admixture analysis provided evidence of four genetic groups that correspond to altitudinal geographic origins and tail morphotypes. Comparative genomic analysis identified targets of selection spanning conserved haplotype structures overlapping genes and gene families relating to hypoxia responses, caudal vertebrae and tail skeleton length, ear morphology, and tail fat-depot structure. Our findings provide novel insights underpinning variation and response to human selection and environmental adaptation, and possible pleiotropic gene interactions in indigenous African sheep genomes, which guaranteed the successful establishment of the species on the continent.

## Introduction

During their human-mediated dispersal across the globe, domestic sheep, *Ovis aries*, encountered and had to adapt to a wide range of novel agro-ecological environments. Natural and artificial selection could have shaped the genomes of domestic sheep to enhance their adaptive fitness to novel African environments. Tail fat deposition could have facilitated such adaptations, and based on tail fat-depot structures, three classes of sheep have been recognized in the continent: fat-tailed, thin-tailed and fat-rumped. Fat-tailed sheep occur in northern, eastern and southern Africa, thin-tailed sheep are predominantly found across western Africa, in Sudan and in parts of northern Africa, and the fat-rumped sheep are restricted to the Horn of Africa (Porter, 2020). Archaeological findings from northern Africa suggest that thin-tailed sheep are the most ancient in the continent and that domestic sheep entered

Africa from the Levant through the Isthmus of Suez in multiple waves (Clutton-Brock, 1993; Marshall, 2000; Gautier, 2006). However, today the fat-tailed sheep predominate in northern Africa. In eastern Africa, the earliest evidence of sheep dates to 4500-3500 BP with rock paintings from the Lake Turkana basin depicting thin-tailed sheep (Barthelme, 1985; MacDonald and MacDonald, 2000), which today are mainly found in Ethiopia (Benishangul-Gumuz and North Gondar regions). Rock paintings from eastern Ethiopian highlands show fat-tailed sheep alongside humpless cattle (Clark and Williams, 1978). The latter arrived on the continent earlier than their humped counterparts which today predominate this region (Gifford-Gonzalez and Hanotte, 2011). Depictions of fat-rumped sheep lack in archaeological findings, thus their origin, entry point and time of diffusion into the continent remains unknown.

Today, Africa is home to ∼383 million sheep (FAOSTAT 2020; accessed July 2020) distributed among 140 phenotypically diverse populations (DAD-IS 2020; accessed July 2020). The continent exhibits marked differences in agro-eco-climates and biophysical challenges (https://iiasa.ac.at/web/home/research/researchPrograms/water/GAEZ_v4.html). It is also home to an exceptional human ethnic agro-pastoral diversity of ancient origin. Together with natural selection for adaptation to diverse environments, human driven selection for economic, socio-cultural and aesthetic traits have shaped the genomes of African livestock, resulting in unprecedented variation within and between populations (e.g., Kim et al., (2020); Gheyas et al., (2021)). The genetic control underpinning such adaptations and variations remain largely unknown. Here, we generated and analysed whole- genome sequences alongside tail morphometric data of indigenous African sheep from Libya, Sudan and Ethiopia. The studied populations represented fat-tailed, thin-tailed and fat-rumped African sheep tail morphotypes and were sampled from different agro-ecologies, ranging from cold high-altitude highlands to hot arid lowland environments. Using the data, we investigated genome-wide levels of genetic variation and structure and used the information to infer demographic-adaptive dynamics of the species in the continent.

## Materials and Methods

### Sample collection

Tissue samples were collected from 150 individuals of 15 indigenous African fat-tailed, fat-rumped and thin-tailed sheep from Ethiopia, Libya and Sudan (Table S1, S2; Figure S1). Genomic DNA was isolated with DNeasy Blood and Tissue Kit (Qiagen) and quality checked with the Nanodrop. Using the Covaris System, 3 μg of genomic DNA were randomly sheared to generate inserts of ∼300 bp. The sheared DNA was end-repaired, A-tailed, adaptor ligated, and amplified with the TruSeq DNA Sample Preparation Kit (Illumina, San Diego, CA, USA). Paired-end sequencing was conducted with the Illumina HiSeq2000. We performed sequence quality checks with the fastQC software (http://www.bioinformatics.bbsrc.ac.uk/projects/fastqc/). The paired-end sequence reads were mapped against the Oar_v3.1.75 sheep reference genome using Burrows-Wheeler Aligner software (Li and Durbin, 2009). We used default parameters (except the “--no-mixed” option) to suppress unpaired alignments. Potential PCR duplicates were filtered using the “REMOVE_DUPLICATES = true” option in “MarkDuplicates” command-line of Picard (http://broadinstitute.github.io/picard). SAMtools (Li and Durbin, 2009) was used to create the index files for reference and bam files. The genome analysis toolkit (GATK) 3.1 (https://gatk.broadinstitute.org) was used to realign the reads to correct for any misalignments arising from the presence of indels (“RealignerTargetCreator” and “IndelRealigner” arguments). “UnifiedGenotyper” and “SelectVariants” of GATK were used to call SNPs. To filter variants and to avoid possible false positives, SNPs with: (i) a phred-scaled quality score < 30, (ii) MQ0 (mapping quality zero) > 4 and quality depth (unfiltered depth of non-reference samples) < 5, and (iii) SNPs with FS (phred-scaled *P* value using Fisher’s exact test) > 200, were filtered out. Following these filtration steps, the retained SNPs were used for analysis.

**Table 1.**
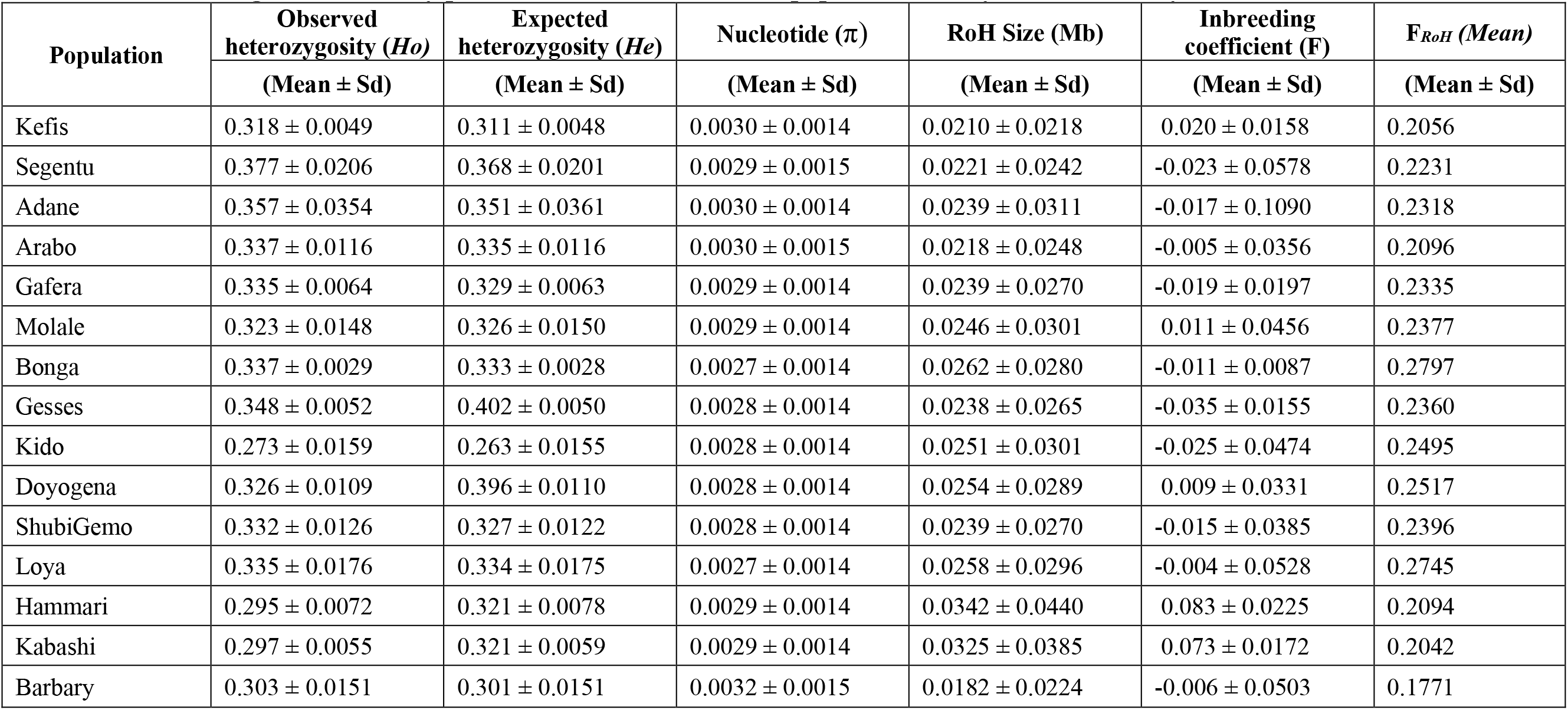
Estimates of genetic diversity parameters for each of the 15 populations analyzed in this study.

### Preparation and measurement of caudal vertebrae

We first subdivided the 15 populations into five groups based on the physical assessment of the tail length and size of fat-depots, *viz*, i) Ethiopian fat-rumped (Kefis, Adane, Arabo, Segentu), ii) Ethiopian long fat-tailed (Bonga, Kido, Gesses, Loya, Shubin Gemo, Doyogena), iii) Ethiopian short fat-tailed (Molale, Gafera) sheep, iv) Sudan long thin-tailed sheep (Hammari, Kabashi) and v) Libya long fat- tailed sheep (Barbary).

One mature animal (≥ 3 permanent pairs of incisor teeth) of Kefis (fat-rumped), Loya (long fat-tailed), Menz (short fat-tailed), Kabashi (long thin-tailed) and Barbary (Libyan long fat-tailed) was selected and slaughtered, the tail eviscerated from the body and the full tail skeleton processed following Zhi et al. (2018). The tail was deskinned, and the flesh and fat gently scrapped from the caudal vertebrae (CVs). The CVs were then incubated in ethanol for five days after which they were cleaned by soaking for two days in 0.5% NaOH. Residual flesh and fat were removed by further soaking the CVs in petrol for three days. The number of CVs were counted and the length (cm) of each CV was determined with a Vernier calliper. The length of the complete tail skeleton was determined with a tape measure. The average length and standard deviation of the CV were calculated with Excel.

### Genetic diversity

The VCFtools v.0.1.15 (Danecek et al., 2011) was used to estimate the observed (*HO*) and expected (*HE*) heterozygosity, and nucleotide diversity (*π*) as indicators of intra-population diversity, while the inbreeding coefficient (*F*) and runs of homozygosity (*RoH*) were estimated as indicators of genome- wide autozygosity. The *RoH* was estimated with the BCFtools software package (Narasimhan et al., 2016). We estimated the *RoH*-derived genomic inbreeding coefficient (FRoH) following McQuillan et al., (2008) as:

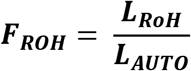

Where *LROH* is the total length of RoH of each individual in the genome and *LAUTO* is the length of the sheep autosomal genome (∼2600 Mb).

### Phylogenetic inference and genetic structure analysis

We visualised population structure using principal component analysis (PCA) performed with Plink 1.9 (Purcell et al., 2007). The first two eigenvectors were plotted to summarise individual relationships. For this analysis, we used 17 million autosomal SNPs, which were retained after filtering out SNPs with minor allele frequency (MAF) < 0.01.

We investigated the proportion of the genome arising from common ancestry using the ADMIXTURE software v.1.3 (Alexander et al., 2009). For this analysis, we used 2.4 million autosomal SNPs that were retained after pruning the 17 million SNPs used in PCA, for linkage disequilibrium (LD). We implemented the block relaxation algorithm with the Kinship (*K*) values set from 2 to 15; five runs were performed for each *K*. A five-fold cross-validation (CV) procedure was used to determine the optimal number of ancestral genomes (*K*) and the proportion of admixture. The PCA and ADMIXTURE results were visualized with GENESIS (Buchmann and Hazelhurst, 2014).

Patterns of population splits and gene flow were assessed with the maximum likelihood tree-based procedure in TreeMix (Pickrell and Pritchard, 2012). The thin-tailed sheep from Sudan were defined as the root. The optimal number of migration events (m) were determined following Pickrell and Pritchard (2012) and the models were visualized with R using the script provided in TreeMix. The contribution of each migration vector that was added to the tree was determined using the “*ƒ* index” (Pickrell and Pritchard, 2012). We modelled the optimal number of migration events and edges (M) from 1 to 10 with the default settings of OptM package in R (https://www.r-pkg.org/pkg/OptM; accessed April 2020).

Five runs were performed for each “M”. The ML phylogenetic tree depicting relationships and the optimal migration event(s) was generated using the default "Evanno"-like method in R.

### Demographic history and dynamics

We estimated the LD parameter (*r^2^*) with Plink v1.9 considering the genetic groups/structure inferred from the phylogeny and ADMIXTURE analysis. The *r^2^* values were sorted and binned in 50 Kb to 10 Mb inter-SNP distances and the genome-wide LD decay plotted using R.

The estimated values of *r^2^* were used to model changes in effective population sizes (*Ne*) over generation time (up to 1000 generations ago) using SNeP (Barbato et al., 2015), for each genetic group/structure inferred by the phylogenetic and ADMIXTURE analysis. The *Ne* values were estimated following Sved (1971) as:

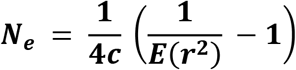

where *c* is the genetic distance in centimorgans (considering 1 cM = 1 Mb), and *E*(*r^2^*) represents the expected LD (*r^2^*) for a distance of *c* Morgan’s. The time intervals denoting the number of generations in the past (*t*) were estimated using *t* = 1/2c (Hayes et al., 2003).

### Genome-wide signatures of selection

We assessed genome-wide signatures of selection for i) adaptation to contrasting environments (e.g., high altitude humid and low altitude dry/arid environments), ii) differences in tail length, and iii) differences in the size of tail fat-depots.

Based on the altitude and agroecology of their geographic home range (Table S2), we classified the study populations into three groups. These were i) sheep from high altitude humid environment (Bonga, Kido, Gesses, Loya, Shubi Gemo, Doyogena and Gafera from Ethiopia), ii) sheep from low altitude hot- dry desert environment (Hammari and Kabashi from Sudan), and iii) sheep from coastal hot-humid arid environment (the Barbary from Libya). Contrasting these three groups allowed for the detection of genomic signatures for adaptation to contrasting environments.

**Table 2.** Candidate regions and genes associated with adaptation to diverse environments as detected in our study.

| Region (base pairs) | Chromosome | Genes | Method | Function |
| --- | --- | --- | --- | --- |
| 253550001-253730000 | 1 | *TF, RAB6B, SRPRB* | $F_{ST}$ | Associated with haemoglobin levels in Tibetans (Simonson et al., 2010) |
| 232750001-232850000 | 2 | *DIS3L2* | $F_{ST}$, $XP-EHH$ | Associated with height variation (de Simoni Gouveia et al., 2017) and thermoregulation (Mcmanus et al. 2011). |
| 77920001-78110000 | 3 | *EPAS1, PRKCE* | $XP-EHH$ | Regulating cellular responses to hypoxia (Frede and Fandrey, 2014) |
| 154130001-154350000 | 3 | *MSRB3* | $H_p$, $F_{ST}$ | high altitude adaptation in dogs (Gou et al., 2014) and Tibetan sheep (Wei et al., 2016). |
| 55940001-56140000 | 7 | *GLDN, CYP19* | $H_p$, $XP-EHH$ | Regulation of male and female reproduction in sheep (Zsolnai et al., 2002). |
| 28560001- 29540000 | 10 | *RXFP2, PDS5B, N4BP2L2* | $F_{ST}$, $XP-EHH$ | implicate its dual role in the development of horns for thermoregulation and enhancing reproductive (Hoefs, 2000; Kardos et al., 2015). |
| 18250001-18490000 | 11 | *NF1, EVI2A, EVI2B, OMG* | $F_{ST}$, $XP-EHH$ | Adaptive response to physical exhaustion (Mwacharo et al., 2017). |
| 27040001- 27180000 | 11 | *KCNAB3, CNTROB, DNAH2, NAA38, KDM6B, TMEM88, CYB5D1, CHD3, RNF227, TRAPPC1* | $H_p$, $F_{ST}$ | Correlated with cognitive performance under chronic stress (Jung et al., 2017) and is down-regulated in response to acute and chronic stress in mice (Terenina et al., 2019). |
| 42670001- 42800000 | 13 | *CHRNA4, SRMS, PTK6, EEF1A2, KCNQ2, ARFGAP1* | $XP-EHH$ | Response to hypoxia (Storz and Moriyama, 2008; Tissot Van Patot and Gassmann, 2011) |
| 69380001- 69520000 | 13 | *PLCG1, ZHX3* | $H_p$ | Response to hypoxia (Storz and Moriyama, 2008; Tissot Van Patot and Gassmann, 2011) |
| 34380001 - 34590000 | 14 | *HSD11B2, ATP6V0D1, AGRP, RIPOR1, CTCF, CARMIL2, ACD, PARD6A, ENKD1, C16orf86, GFOD2* | $F_{ST}$ | Associated with salt sensitivity (Alikhani-Koupaei et al., 2007) |
| 35110001-35280000 | 15 | *PLEKHA7, C11orf58* | $F_{ST}$ | Associated with salt sensitivity (Lin et al., 2011; Taal et al., 2012) |
| 17230001-17480000 | 20 | *VEGFA, MAD2L1BP, RSPH9, MRPS18A* | $F_{ST}$ | Cerebrovascular adaptation to chronic hypoxia in sheep (Castillo-Melendez et al., 2015) |
| 4050001- 4170000 | 25 | *EGLN1, GNPAT, EXOC8, SPRTN* | $H_p$, $F_{ST}$, $XP-EHH$ | Regulating cellular responses to hypoxia (Frede and Fandrey, 2014) |

Based on the average lengths of the CVs and the length of the complete tail skeleton, we classified the studied populations into two groups, i) short-tailed sheep comprising short fat-tailed and fat-rumped sheep, and ii) long-tailed sheep comprising long fat-tailed and long thin-tailed sheep. These were used to investigate genomic regions associated with differences in CV sizes and in the length of the tail skeleton.

To identify candidate regions associated with differences in the size of fat tail-depots, we contrasted the long thin-tailed sheep with the long fat-tailed, fat-rumped and short fat-tailed sheep, respectively.

Signatures of selection were investigated by analysing 34.8 million autosomal SNPs with three methods: i) within-group pooled heterozygosity (*Hp*) (Rubin et al., 2010), ii) genetic differentiation based on *F*ST (Weir and Cockerham, 1984), and iii) the haplotype-based *XP–EHH* (Sabeti et al., 2007). A sliding window was used to perform the *Hp* and *FST* analysis with VCFtools v.0.1.15. Based on LD decay trend, a 100 kb non-overlapping window was chosen for the analysis. By exploring several sliding-window distances (10 kb, 20 kb, 50 kb, 75 kb, 100 kb), the 10 kb sliding-window distance gave the best resolution of the selection signals and was thus chosen for our analysis.

We estimated the genome-wide *Hp* statistic using the formula:

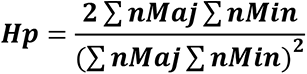

where ∑nMaj and ∑nMin are the sum of the number of the major allele and the number of the minor allele, respectively. We transformed the *Hp* values into Z-scores using the formula:

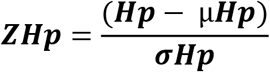

Where *µHp* is the average of the overall heterozygosity and *σHp* is the standard deviation for all the windows within the test group.

The *FST* (Weir and Cockerham, 1984) values were estimated for each SNP in each window between test groups with the formulae:

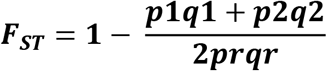

where *p1, p2* and *q1, q2* are the frequencies of alleles “A” and “a” in the first and second test groups, respectively, and *pr* and *qr* are the frequencies of alleles “A” and “a”, respectively, across the test groups. The *FST* values were standardized into *Z*-scores as follows:

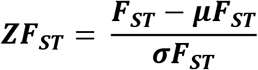

where *μFST* is the overall average value of *FST* and *σFST* is the standard deviation derived from all the windows tested between test groups.

The *XP-EHH* method contrasts extended haplotype homozygosity (*EHH*) between populations in detecting selection signatures (Sabeti et al., 2007). The approach estimates and contrasts the *iES* statistic (the pattern of integrated *EHH* of the same allele) between populations as follows:

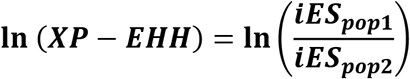

Where *XP − EHH* is the cross-population differentiation, *iES*_*pop*1_ is the integrated *EHHS* for the targeted sheep group and *iES*_*pop*2_ is the integrated *EHHS* for the reference sheep group.

For the analysis, haplotype phasing was inferred on all bi-allelic SNPs with BEAGLE (Browning and Browning, 2007). Assuming 100 Mb = 1 Morgan (Quinlan and Hall, 2010), phased haplotypes were converted to IMPUTE format using VCFtools v.0.1.15. The generated haplotypes were then used to estimate *XP-EHH* scores in pair-wise comparisons between test groups using HAPBIN (Maclean et al., 2015). To determine the empirical significance of the *XP-EHH* statistic, *XP-EHH* scores were normalized by subtracting the mean and dividing it with the standard deviation of all the scores. Negative *XP-EHH* scores indicate selection in the reference group, while positive scores indicate selection in the test group.

The top negative 99% values for *ZHp*, and the top 0.1% of the empirical distributions of the *ZF*ST and *XP-EHH* scores were used as the cut-off threshold to detect outliers and define candidate regions under selection. For a region to be considered a candidate under selection, it had to span at least three significant SNPs. Candidate regions that overlapped between at least two methods were identified and merged using Bedtools v.2.25.0 (Quinlan and Hall, 2010).

### Haplotype structure analysis

We further investigated the candidate region intervals and putative genes for evidence of conserved haplotype structures in a test group. We estimated haplotype frequencies and visualized the haplotype structures using the *hapFLK* software (Fariello et al., 2013) employing the scripts from the *hapFLK* homepage (https://forge-dga.jouy.inra.fr/projects/hapflk).

### Functional annotation

The candidate regions were annotated based on the Oar-v3.1.75 sheep reference genome assembly with the *Ensembl BioMart* tool (http://www.ensembl.org/biomart). Using the annotated genes found in all the candidate regions, DAVID v6.8 (Huang et al., 2009) and KOBAS v3.0 (Xie et al., 2011) were used to perform functional enrichment analysis. The *O. aries* annotation was used as the background species. To interpret the gene functions in a livestock context, we retrieved information available on their functional effects from literature.

## Results

### Variant discovery and annotation

The overall alignment rate of the reads to the ovine reference sequence was 98.56% with an average sequencing depth of ∼54x for Ethiopian and Libyan samples and ∼10x for Sudanese samples. The summary statistics for the sequence parameters are shown in Supplementary Table S3. In general, a comparison with the *O. aries* dbSNP database, revealed that ∼6% of the SNPs and InDels were not present in the database (Table S3, S4). Of the 34,857,882 SNPs generated in this study, ∼36.8%, ∼55.9% and ∼0.79% were intronic, intergenic and exonic, respectively (Table S5; Figure S2).

**Table 3.**
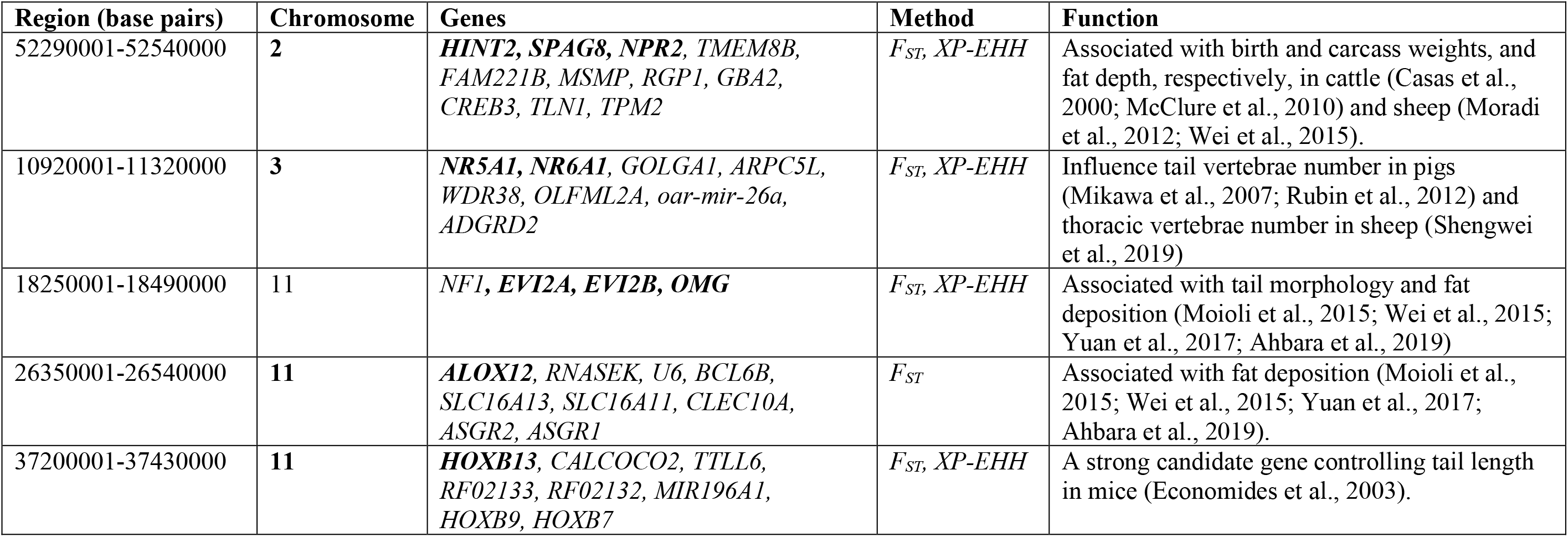
Candidate regions and genes associated with phenotypic traits (tail length and tail fat-depots) as detected in our study.

### Tail skeleton morphometry

The tail and skeletal tail characteristics of Menz, Kefis, Loya, Barbary and Kabashi are shown in Figure 1a. The tail of Loya, a long fat-tailed sheep, extends to the hock joint while that of Menz and Kefis representing short fat-tailed and fat-rumped sheep, respectively does not extend beyond the hock joint. The thin-tailed Kabashi and long fat-tailed Barbary sheep have the longest tails, which extend beyond the hock joint, with the tail of SD sheep almost touching the ground. The short fat-tailed and fat-rumped sheep have 8-10 CVs, the Ethiopian long-tailed has 18, and LB and SD have 22 each. SD has the highest average length of CV (2.72 ± 0.364 cm), followed by the Ethiopian long-tailed (2.42 ± 0.243 cm), fat- rumped (2.24 ± 0.113 cm) and LB (2.20 ± 0.183 cm). Although LB has 22 CV’s, their average lengths are the shortest resulting in a shorter overall tail skeletal length compared to SD. Based on the average lengths of the individual CV’s and the length of the complete tail skeleton, we classified the study populations into two groups, i) long-tail sheep comprising SD and the Ethiopian long-tail and, ii) short- tail sheep comprising LB and the Ethiopian fat-rumped.

**Figure 1.**
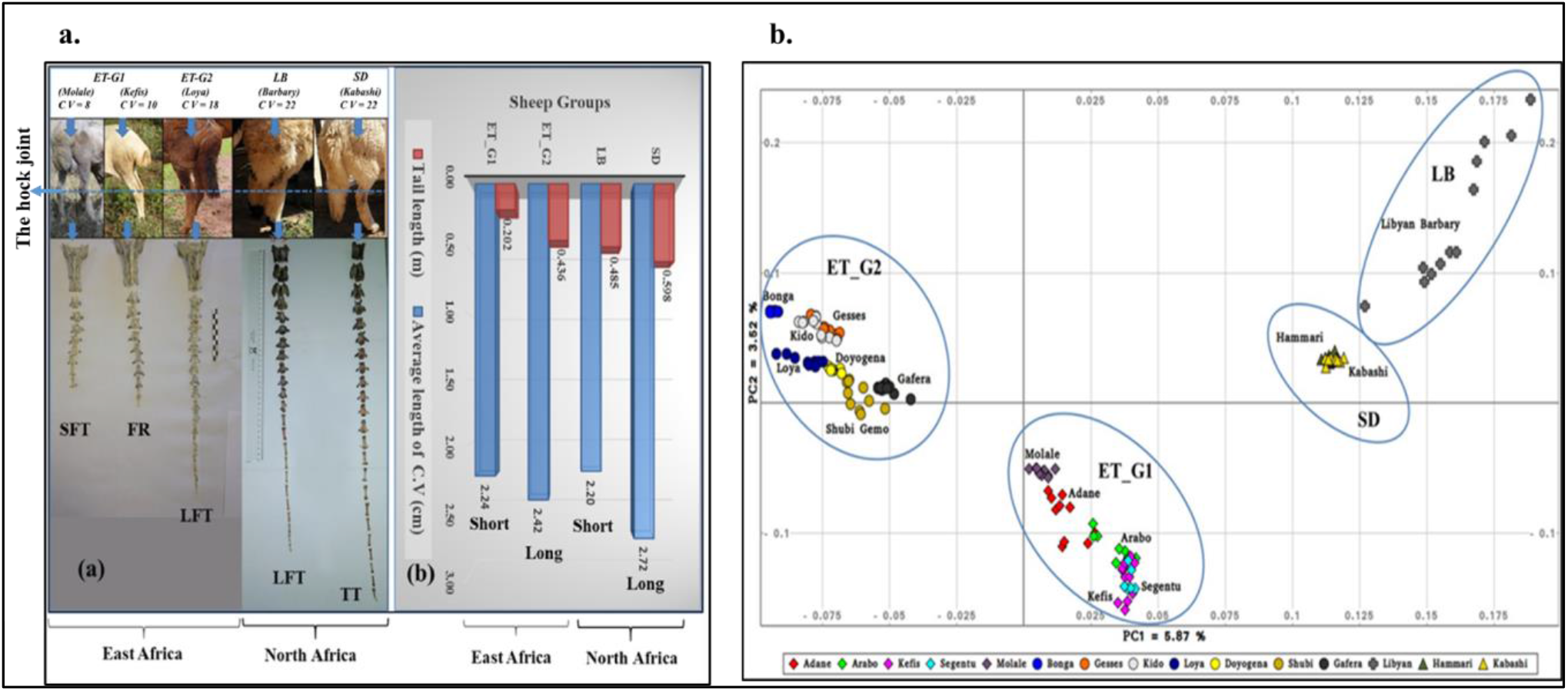
Tail phenotypes of different sheep groups: (a) visual length and number of caudal vertebrae (CV). (b) Tail length (meters) and average length (cm) of single vertebra. SFT = short fat-tailed, FR = Fat-rumped, LFT = Long fat-tailed and TT = Thin-tailed; (c). Clustering pattern of the 15 sheep populations studied as revealed by principal component analysis.

### Genetic diversity

The average estimates of *HO*, *HE*, *π*, *F, RoH* and *FRoH*, and their standard deviations are shown in Table 1. The least and most diverse populations based on *HO* and *HE* estimates were Kido and Segentu, respectively. The highest value of *π* was in Barbary and the lowest in Bonga and Loya. Excepting Hammari, Kabashi and Menz, the other populations showed low values of inbreeding (*F* < 1.0%). Hammari and Kabashi reported the largest mean sizes of *RoH* and Barbary had the smallest (Table 1; Figure S3). Barbary also had the lowest *FRoH*, while Bonga and Loya had the highest (Table 1).

### Population structure

The PCA (Figure 1b) separated Ethiopian sheep into two genetic groups corresponding to the ones reported by Ahbara et al. (2019) based on the analysis of 50K SNP genotypes. For consistency, we adopted Ahbara et al. (2019) nomenclature viz: “ET-G1” comprising fat-rumped sheep (Kefis, Adane, Arabo, Segentu) and the short fat-tailed Molale/Menz sheep)), and “ET-G2” comprising long fat-tailed sheep (Bonga, Kido, Gesses, Loya, Shubi Gemo, Doyogena) and the short fat-tailed Gafera sheep)). The thin-tailed (Hammari and Kabashi) sheep from Sudan (SD) and the fat-tailed Barbary (LB) sheep from Libya clustered close but separate from each other and from Ethiopian sheep. The Barbary sheep comprise a more genetically diverse group than the other populations.

To determine the optimal clustering pattern generated by ADMIXTURE, we calculated and plotted the CV error for each *K*. The lowest CV error occurred at *K* = 3 (Figure S4a). The distribution of the three genetic backgrounds, named here A, B and C, among the 15 study populations is shown in Figure S4b. The “A” and “B” backgrounds predominated in ET-G1 and ET-G2 populations, respectively. The “C” background occurred in SD and LB. Interestingly, the “A” and “B” backgrounds were also present in SD and LB but, the “C” background occurred at very low frequencies in a few individuals of ET-G1 and ET-G2. There was a clear divergence of Molale sheep and roughly 50 % of Gafera and Adane populations whose genomes are defined by the D background at 4≤*K*≤5. Furthermore, at *K*=5, a fifth ancestry (E) is observed that separates the western populations from the remainder of the Ethiopian populations (Figure S4b).

The pattern of population splits revealed by TreeMix mirrors that of PCA (Figure 2a). The long fat-tailed (Gesses, Kido, Bonga, Doyogena, Loya, Shubi Gemo) and the short fat-tailed (Gafera) sheep formed one group. The fat-rumped (Adane, Arabo, Kefis, Segentu) and the short fat-tailed (Menz) sheep formed another group. The Libya Barbary (LB) and the Sudan thin-tailed (SD) sheep represented two separate lineages. TreeMix also provided evidence of gene-flow. The first seven migration vertices accounted for > 90% of the model significance, with the first migration edge accounting for the highest contribution. Migration vectors ≥ 8 resulted in small incremental changes in the value of the model significance. We therefore chose m = 7 as the best predictive value of the optimal migration model.

**Figure 2.**
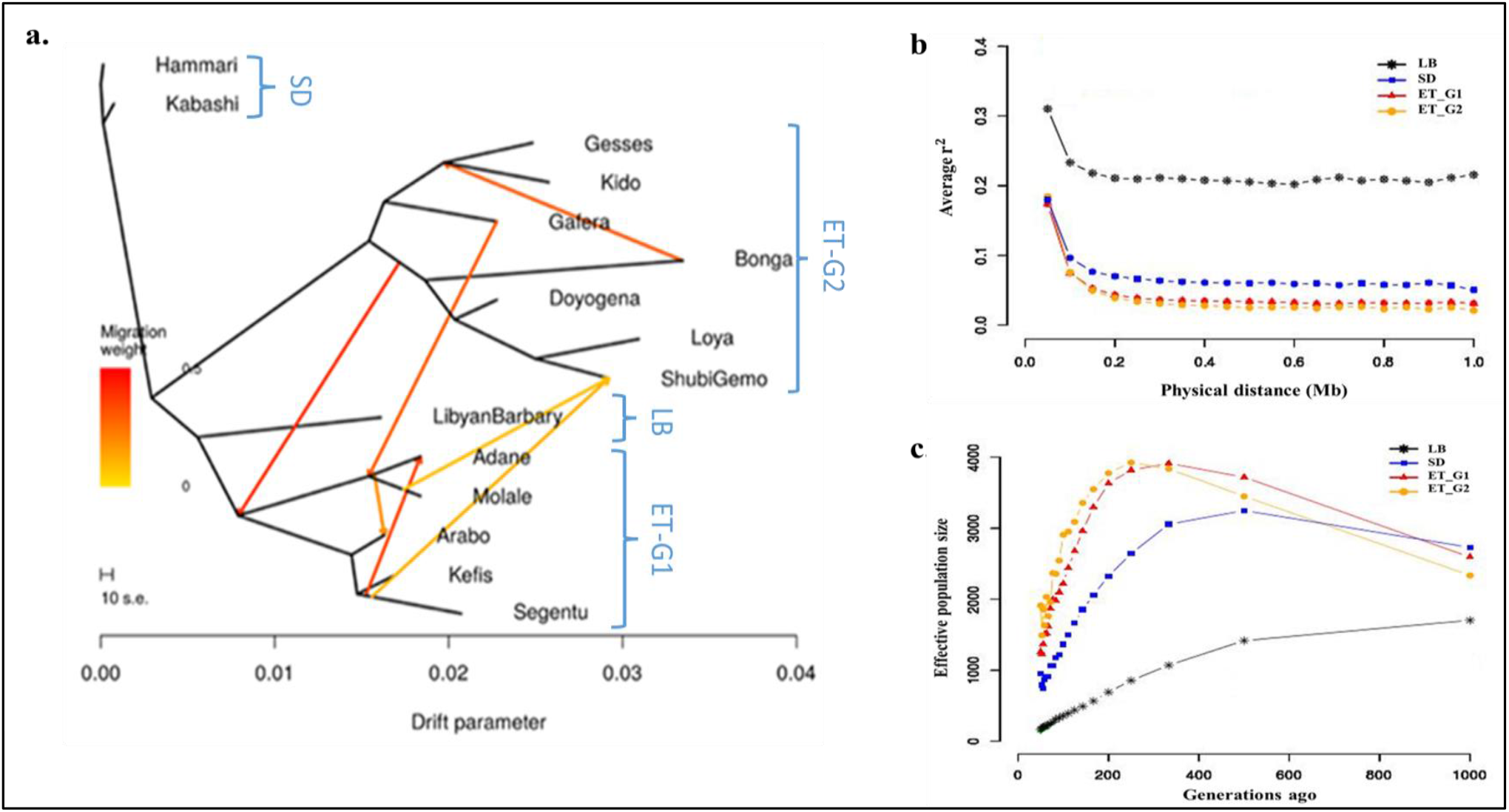
(a). Phylogenetic network inference based on TreeMix analysis depicting the relationships between Ethiopian, Libyan and Sudanese sheep populations. The first seven migration edges are shown with arrows pointing the direction of gene flow toward the recipient group and coloured according to the percentage of ancestry received from the donor, (b). Patterns of linkage disequilibrium (*r^2^*) from 0 to 1Mb for the four sheep groups revealed by PCA and ADMIXTURE analysis and (c). Average estimated effective population size (*Ne*) of the four sheep groups revealed by PCA and ADMIXTURE analysis over the past 1000 generations.

Four strong (migration weight > 0.35) migration events were revealed (Figure 2a). They showed geneflow from i) central and southern Ethiopian (Bonga, Doyogena, Loya, ShubiGemo (long fat-tailed)) sheep to eastern Ethiopian (Adane, Molale, Arabo, Kefis (fat-rumped)) sheep, ii) Bonga to the cluster of Gesses and Kido sheep in the West of the country, iii) Gafera to the cluster of Menz and Adane sheep in Central Ethiopia, and iv) Segentu to Adane. Three weak (migration weight < 0.2) migration events were also revealed; two involved gene flow from Menz and Segentu, respectively to Shubi Gemo, and from the cluster of Adane and Menz to Arabo.

The average values of *r^2^* reflecting genome-wide LD were lowest in ET-G2 and highest in LB (Figure 2b; Table S6). Irrespective of the genetic group, the trend in *r^2^* showed a rapid decline at the 10 kb to 100 kb distance interval. The trends in *Ne* for each genetic group are shown in Figure 2c and Table S7. SD and LB showed the highest and lowest *N*e, respectively at 1000 generations ago. ET-G1 showed higher *N*e than ET-G2 up to 300 generations ago following which the opposite is observed. Except LB whose *N*e declined gradually up to 50 generations ago, that of ET-G1, ET-G2 and SD increased gradually up to 300 generations ago, and then declined rapidly up to 50 generations ago.

### Genome-wide scans for signatures of selection

The PCA revealed four genetic groups (ET-G1, ET-G2, SD and LB), which were supported by TreeMix. Because the north African SD and LB shared a prominent background (C), these four groups were reduced to only three ancestries at the optimal cross validation error value of *K*=3 (Figure S4b). The ET-G2 comprised populations from a high-altitude humid environment while LB and SD comprised sheep from a coastal hot-humid arid environment in Libya and low altitude hot-dry desert environment in Sudan, respectively (Table S1; S2). The ET-G1 included populations from diverse environments.

#### Selection signatures for populations from high-altitude humid regions

We used the ET-G2 group to investigate selection signatures for adaptation to high-altitude African environments (Figure 3). When this group (ET-G2) was contrasted with LB and SD, the three analytical approaches (*Hp, FST, XP-EHH*), either independently or in combination, identified 91 candidate regions spanning 250 genes (Table S8). These genes were used for functional enrichment analysis, and we considered the genes that comprised the three most significant (*P*-value ≤ 0.01) KEGG Pathways and GO terms as the primary candidates driving high altitude adaptation (Table S9). The genes that have been associated with high altitude environmental adaptation and were detected in our candidate selection regions are summarised in Table 2. The top-two most significant GO biological process terms were “response to hypoxia (GO:0001666)” and “positive regulation of angiogenesis (GO:0045766,)”. Genes associated with “response to hypoxia” were *EPAS1, EGLN1, CHRNA4, VEGFA, NF1* and *ADSL*; the ones associated with “positive regulation of angiogenesis” were *VEGFA* and *FGF2*. The top-two most significant KEGG pathways were “tuberculosis (oas05152)” and “HIF-1 signaling (oas04066)”. Genes associated with the former were *IL18, PLK3, TLR1, CATHL3, CALML4, BAC5, TLR6* and *SC5*, and the ones associated with the latter were *TF, EGLN1, VEGFA* and *PLCG1* (Figure S5).

**Figure 3.**
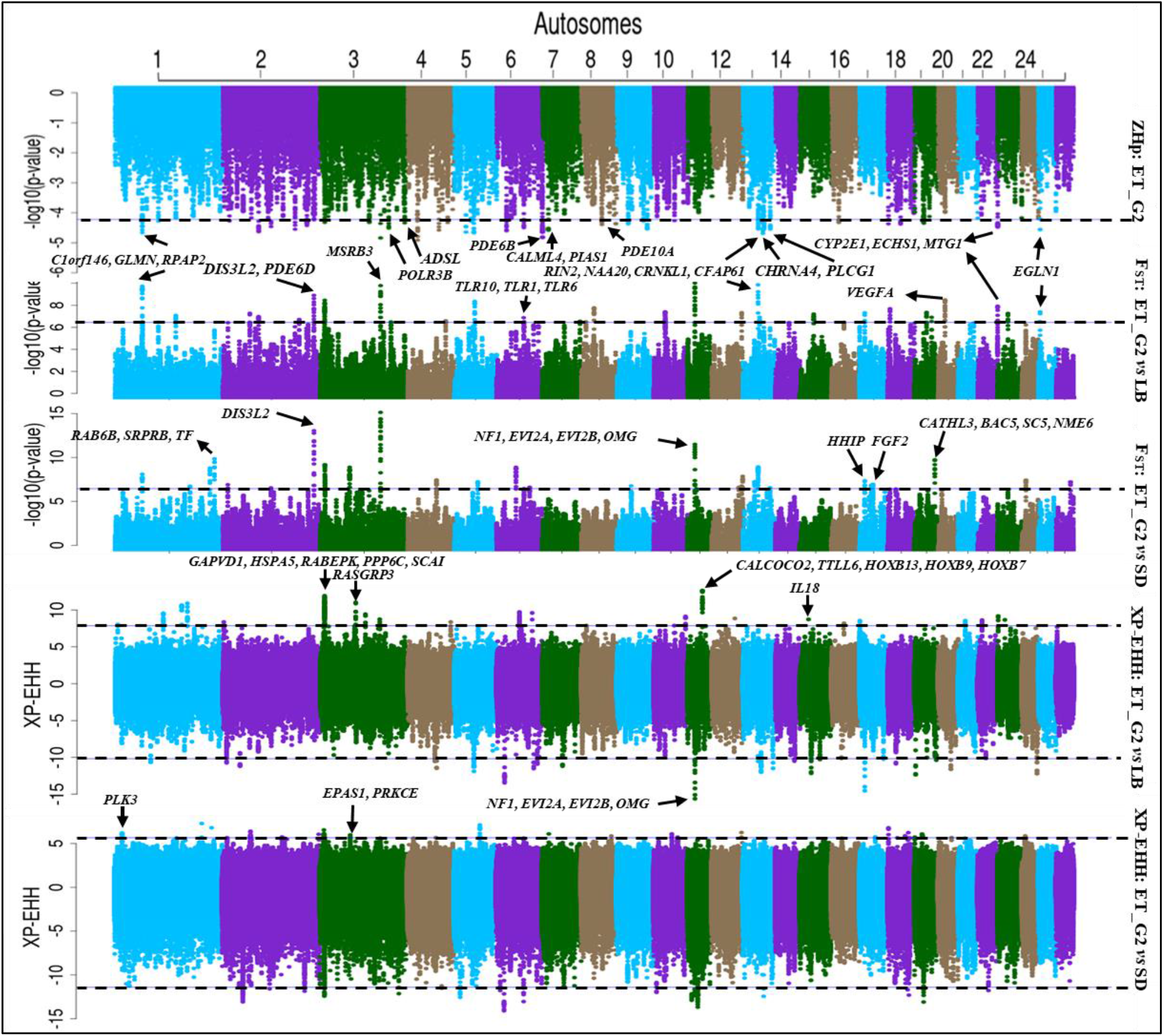
Manhattan plot of genome-wide distribution of ZHp, FST, and XP-EHH values generated from the comparative analysis involving Ethiopian long fat-tailed (ET-G2) sheep with Libyan Barbary (LB) and long thin-tailed (SD) sheep group from Sudan.

We explored haplotype structures around *EGLN1* and *TF* genes (Figure 4a, and 4b) that have been reported in previous studies to be playing a significant role in high altitude adaptation. The analysis revealed the presence of a haplotype that was fixed in ET-G2 (frequency = 1.00), almost fixed in SD (frequency = 0.95) and segregating in LB (frequency = 0.625).

**Figure 4.**
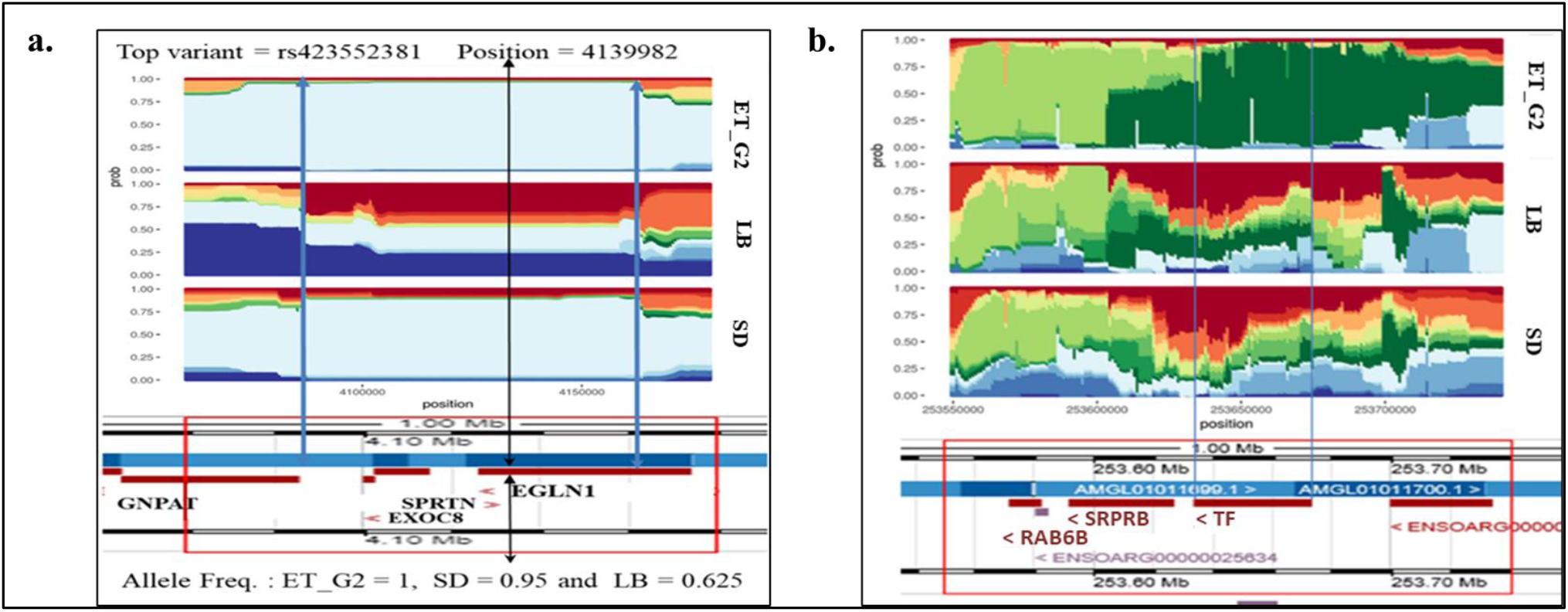
Haplotype structure at the candidate region spanning *EGLN1* on OAR25. The position of the most significant variant is shown by the black arrow, allele frequencies in the different groups are shown below. 4b. Haplotype structure around the candidate region spanning the *TF* gene on OAR1.

#### Selection signatures for low altitude hot-dry desert (Sudan), and coastal hot-humid arid (Libya) environments

We contrasted both LB and SD with ET-G2 to identify selection signatures for adaptation to low-altitude hot-dry desert and coastal hot-humid arid African environments (Figure 5). The three tests revealed 123 candidate regions spanning 182 genes (Table S10). Two of these regions were identified by *Hp* and *FST*, six by *Hp* and *XP-EHH*, three by *XP-EHH* and *FST*, two by all the three methods and one by *XP-EHH*. The two regions identified by all the three methods occurred on OAR6 and OAR11, respectively. The region on OAR6 spanned no genes while the one on OAR11 was amongst the strongest and spanned four genes, *NF1, EVI2A, EVI2B* and *OMG* (Figure 5). Another strong candidate region that was supported by *Hp* and *FST* occurred on OAR2 and spanned *DIS3L2* (Figure 5). The haplotype structure analysis around *DIS3L2* showed a haplotype that was approaching fixation in LB and SD (Figure 6a). Another strong candidate region was found on OAR3 and overlapped *MSRB3* (Figure 5, 6b). The ET-around *MSRB3* compared to LB and SD groups, which are characterised by long drooping ears (Figure 6b). Several LB-specific candidate regions were also revealed; one occurred on OAR7 and spanned *GLDN* and *CYP19* genes. Another was on OAR15 and spanned *PLEKHA7* and *C11orf58*. A distinct haplotype occurred around *PLEKHA7* (Figure 6c). Two other regions spanning distinct haplotypes around *PDS5B* and *RXFP2* genes were found 570 kb apart on OAR10 (Figure 6d). In total, 182 genes found in 123 regions (Table S10) were identified and used for functional enrichment analysis yielding 14 highly significant GO terms and KEGG pathways (Table S11). The top-most highly significant KEGG pathways included “Aldosterone synthesis and secretion (oas04925)” and “Vasopressin- regulated water reabsorption (oas04962)”. The candidate genes associated with low altitude hot-dry desert (Sudan), and coastal hot-humid arid (Libya) environments adaptation are summarised in Table 2.

**Figure 5.**
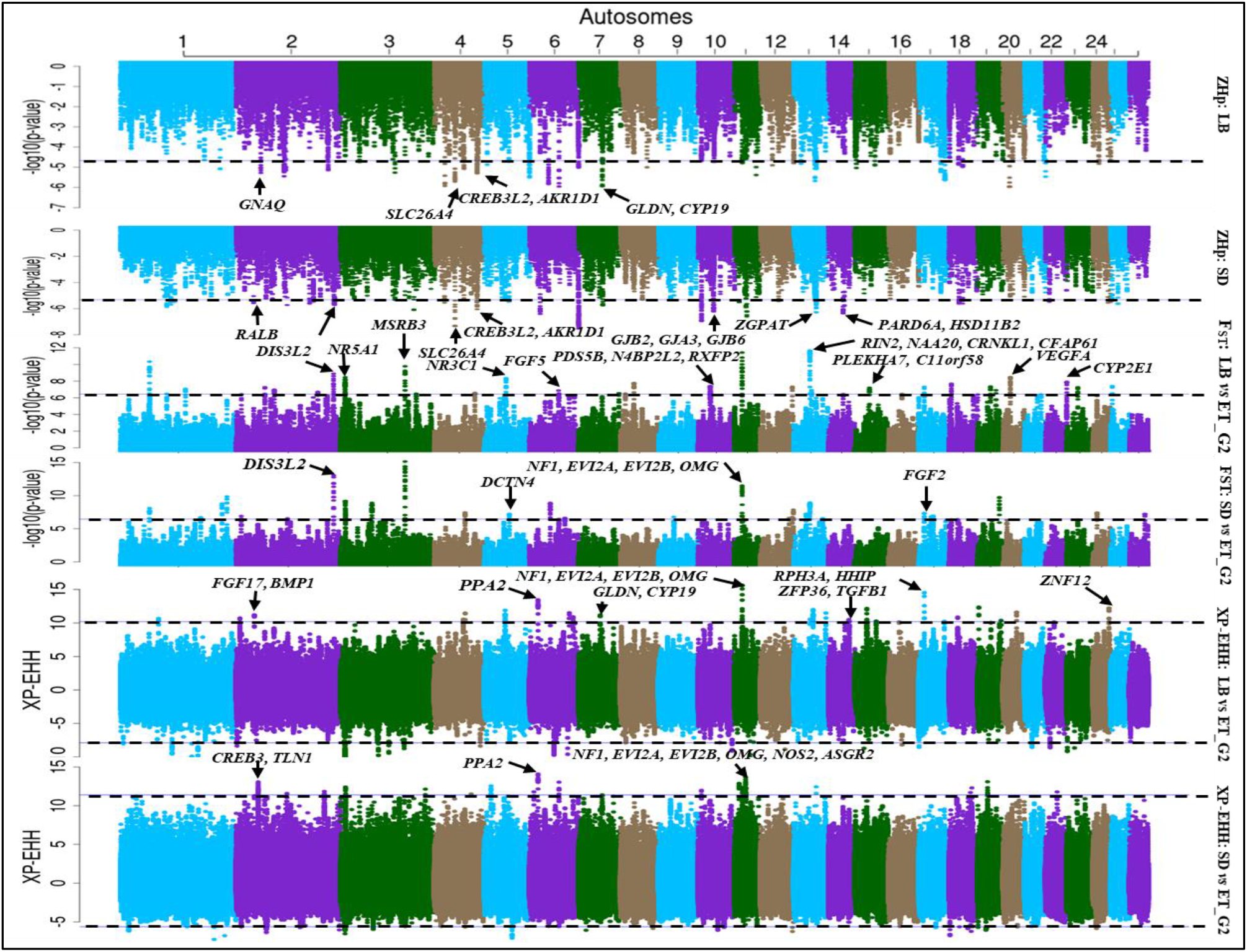
Manhattan plot of genome-wide distribution of SNPs and the candidate regions associated with arid/desert low-altitude environment as detected by ZHp in LB and SD sheep, and detected by FST and XP-EHH by contrasting LB and SD sheep with ET-G2 group from high-altitude African environments.

**Figure 6.**
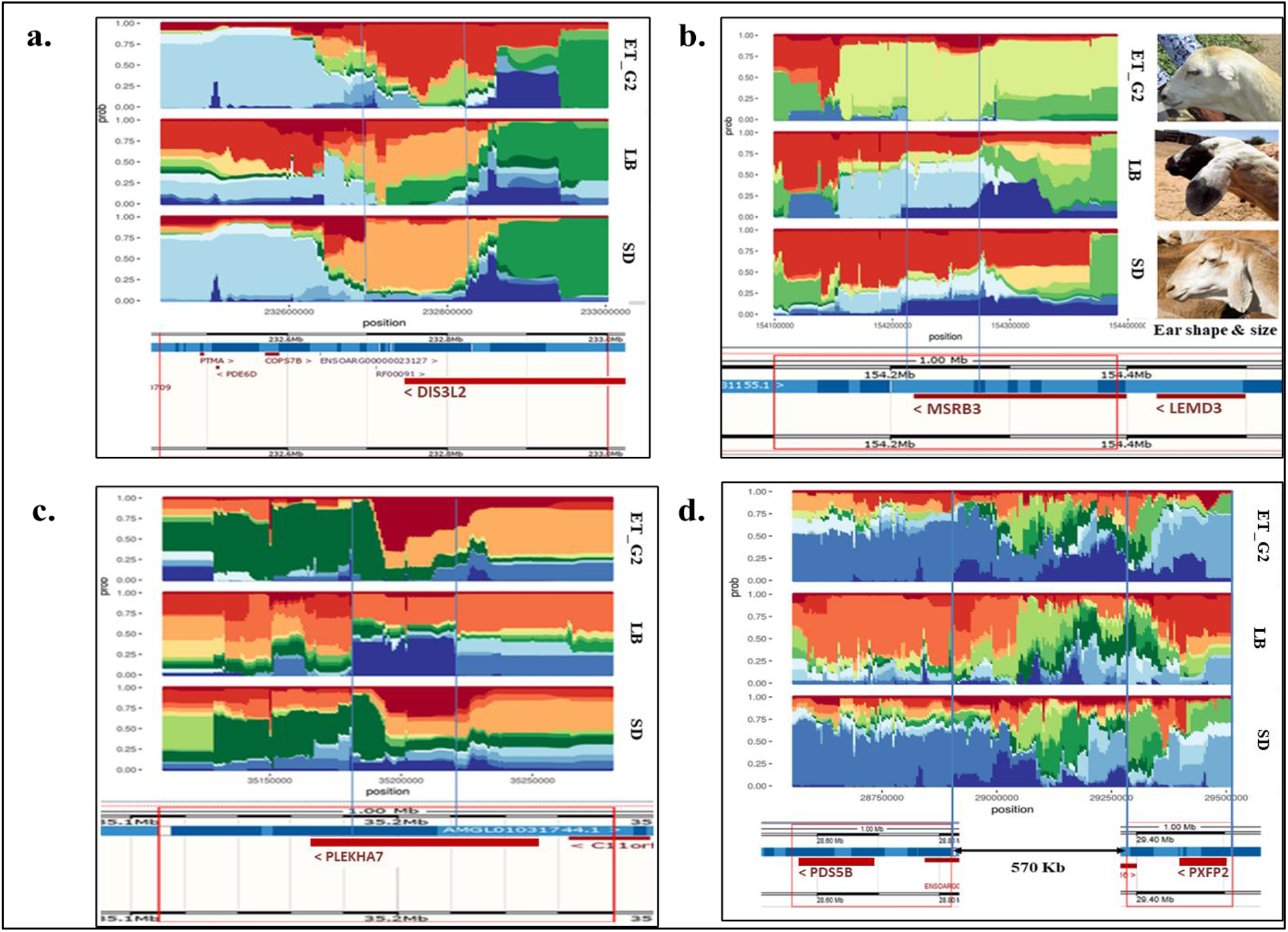
(a). Haplotype structure around the candidate region on OAR2 spanning the *DIS3L2* likely associated with limb length, (b). Haplotype structure around the candidate region on OAR3 spanning *MSRB3* likely associated with ear size and shape, (c). Haplotype structure around the candidate region on OAR15 spanning *PLEKHA7* likely involved in salt-sensitivity, blood pressure, kidney function and hypertension, (d). Haplotype structure around *PDS5B* and *RXFP2* genes likely associated with heat stress and horn characters on OAR10.

#### Signatures of selection for differences in tail length and fat-depot size

To identify candidate regions associated with tail length, we took SD and ET-G2 to represent long-tailed sheep and LB and ET-G1 to represent short-tailed sheep, based on the lengths of the individual CV’s and the complete tail skeleton. We analysed selection signatures with *Hp* (Figure 7), *FST* (Figure 8) and *XP-EHH* (Figure 9, 10) and identified 68 candidate regions across 21 autosomes (Table S12). Two candidate regions were identified by the three tests and 15 regions by at least two tests. The remaining 51 regions were identified by only one test; they included 29 identified by *FST* and 22 by *XP-EHH*. The annotation of these candidate regions identified 218 genes (Table S12). The top-most tail-formation related candidate genes are shown in Table 3. The first region identified by all the three tests occurred on OAR3 and spanned six genes (*GAPVD1, HSPA5, RABEPK, PPP6C, SCAI, ENSOARG00000025028*). The second occurred on OAR6 and spanned no genes. One strong candidate region that overlapped eight genes (*CALCOCO2, TTLL6, HOXB13, RF02133, RF02132, MIR196A1, HOXB9, HOXB7*) was identified by *FST* and *XP-EHH* on OAR11 (Table 3; Figure 8, 9 and 10). Of these eight genes, *HOXB13* was identified as a strong candidate gene controlling tail length in mice (Economides et al., 2003). An analysis of haplotype structures around this gene (Figure 11) revealed a conserved haplotype with a frequency of 0.843 and 0.75 in ET-G2 and LB, respectively. A second strong candidate region overlapping seven genes (*GOLGA1, ARPC5L, WDR38, OLFML2A, NR5A1, NR6A1, ADGRD2*) occurred on OAR3 (Table 3; Figure 8, 9 and 10). Two of the genes (*NR5A1* and *NR6A1*) have been associated with vertebrae number (Yang et al., 2009).

**Figure 7.**
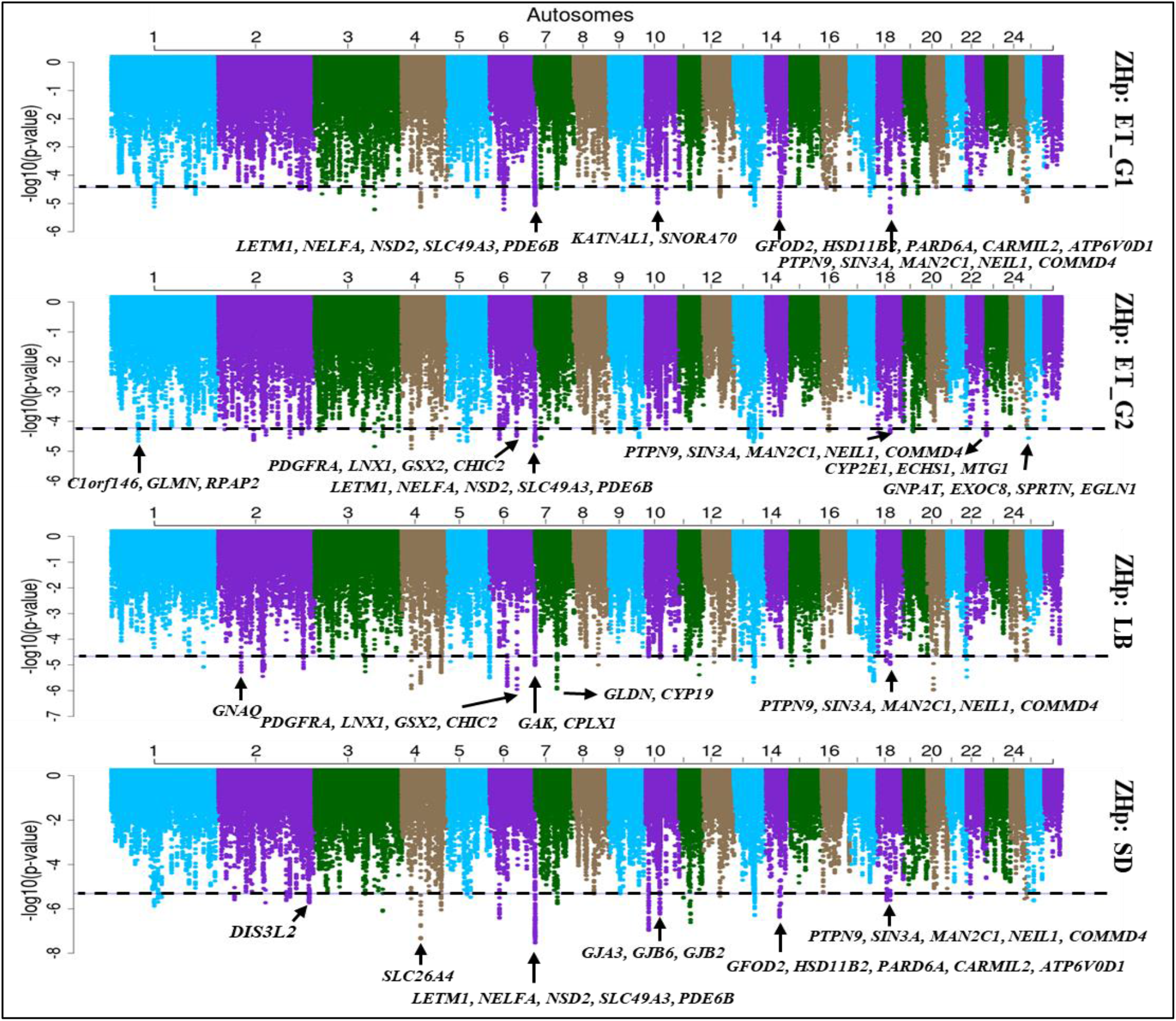
Manhattan plot of genome-wide distribution of ZHp values in Ethiopian fat-rumped (ET-G1), long fat-tailed (ET-G2), Libyan Barbary (LB) and Sudanese thin-tailed (SD) sheep group.

**Figure 8.**
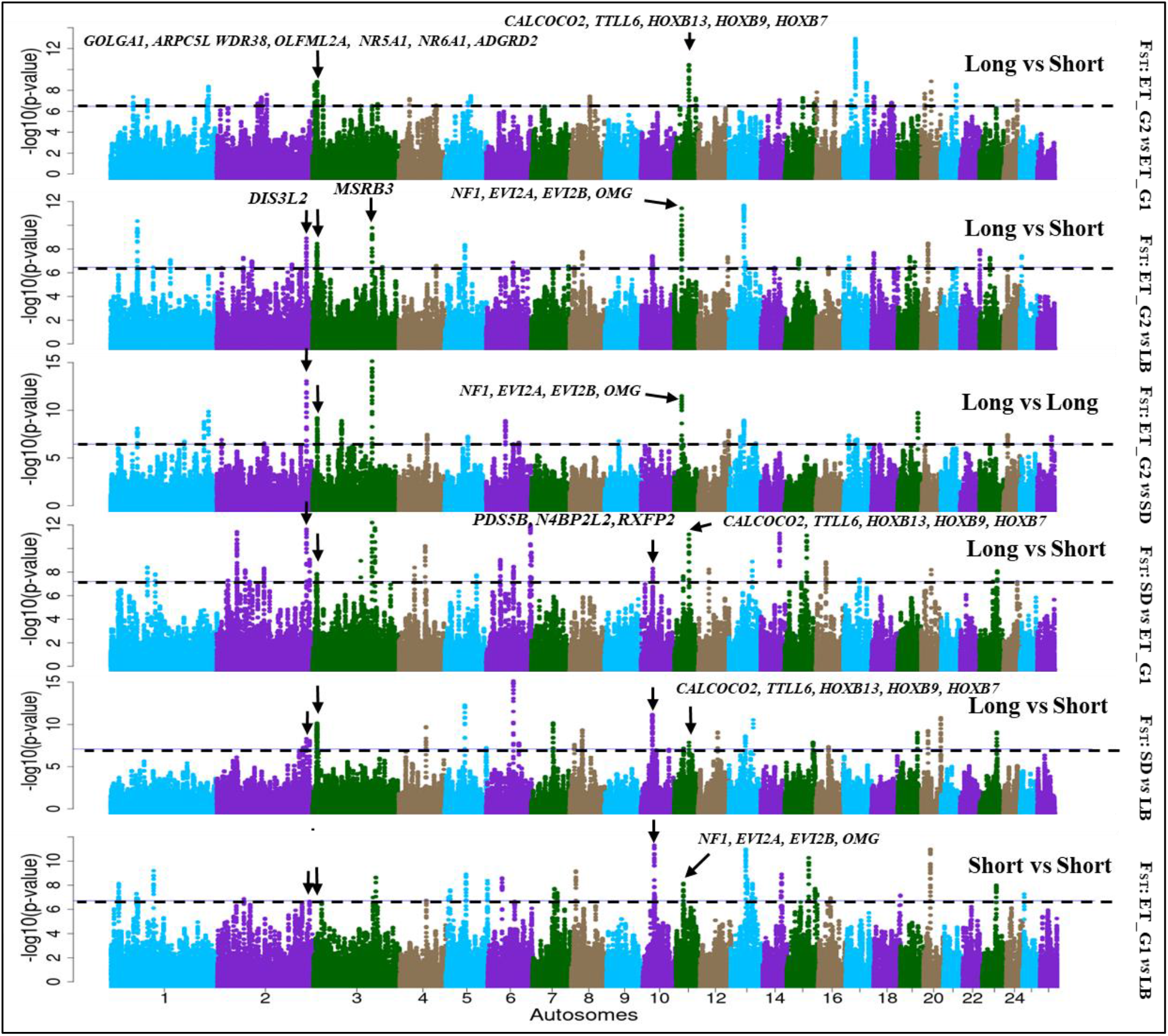
Manhattan plot showing genome-wide distribution of ZFST values following the pairwise comparative analysis between long-tailed Sudan sheep (SD), long-tailed Ethiopian (ET-G2), short-tailed Libyan Barbary (LB) and short-tailed Ethiopian (ET-G1) sheep groups.

**Figure 9.**
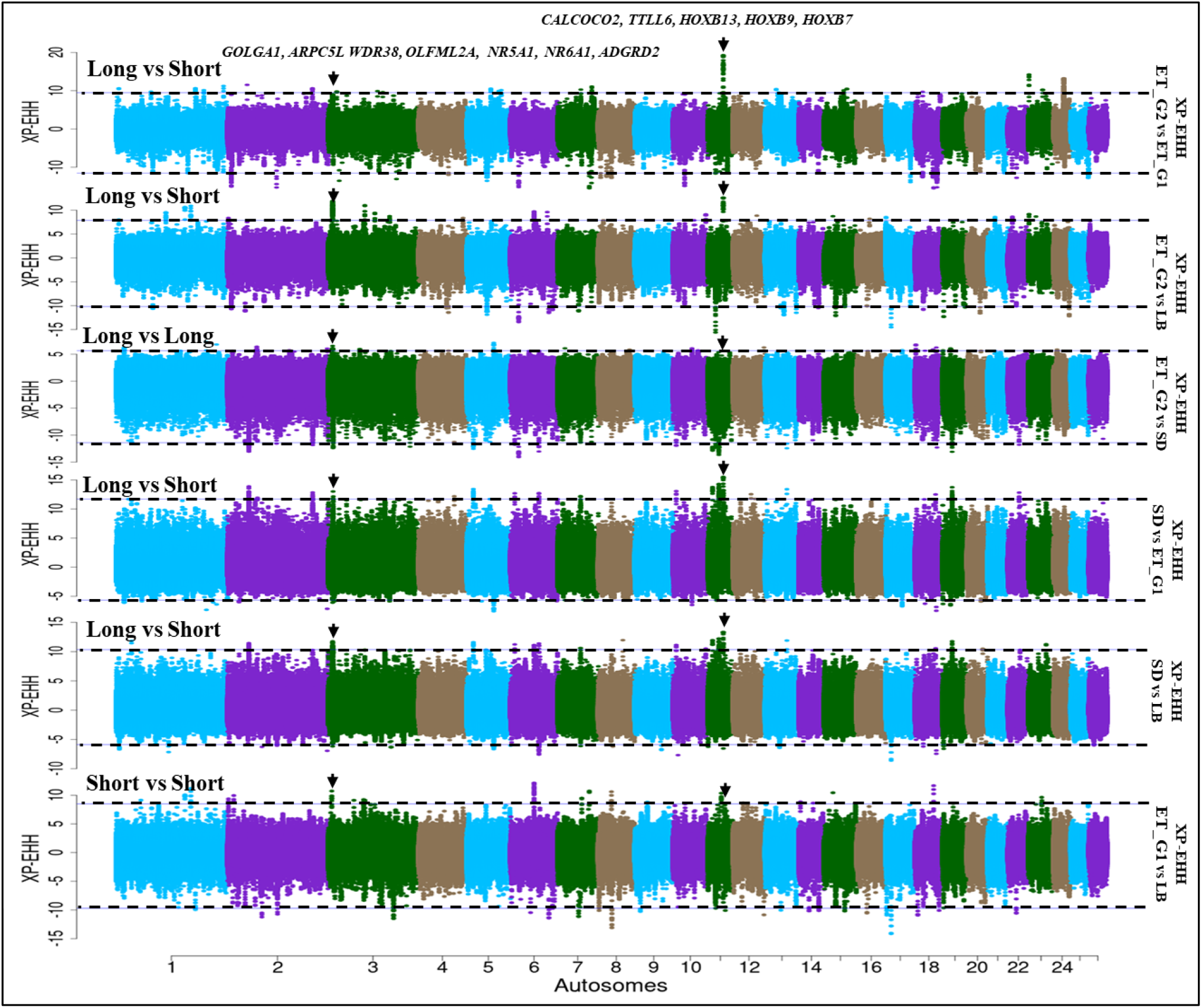
Manhattan plot showing genome-wide distribution of XP-EHH values following pairwise comparative analysis between long-tailed Sudan sheep (SD), long-tailed Ethiopian (ET-G2), short-tailed Libyan Barbary (LB) and short-tailed Ethiopian (ET-G1) sheep groups.

**Figure 10.**
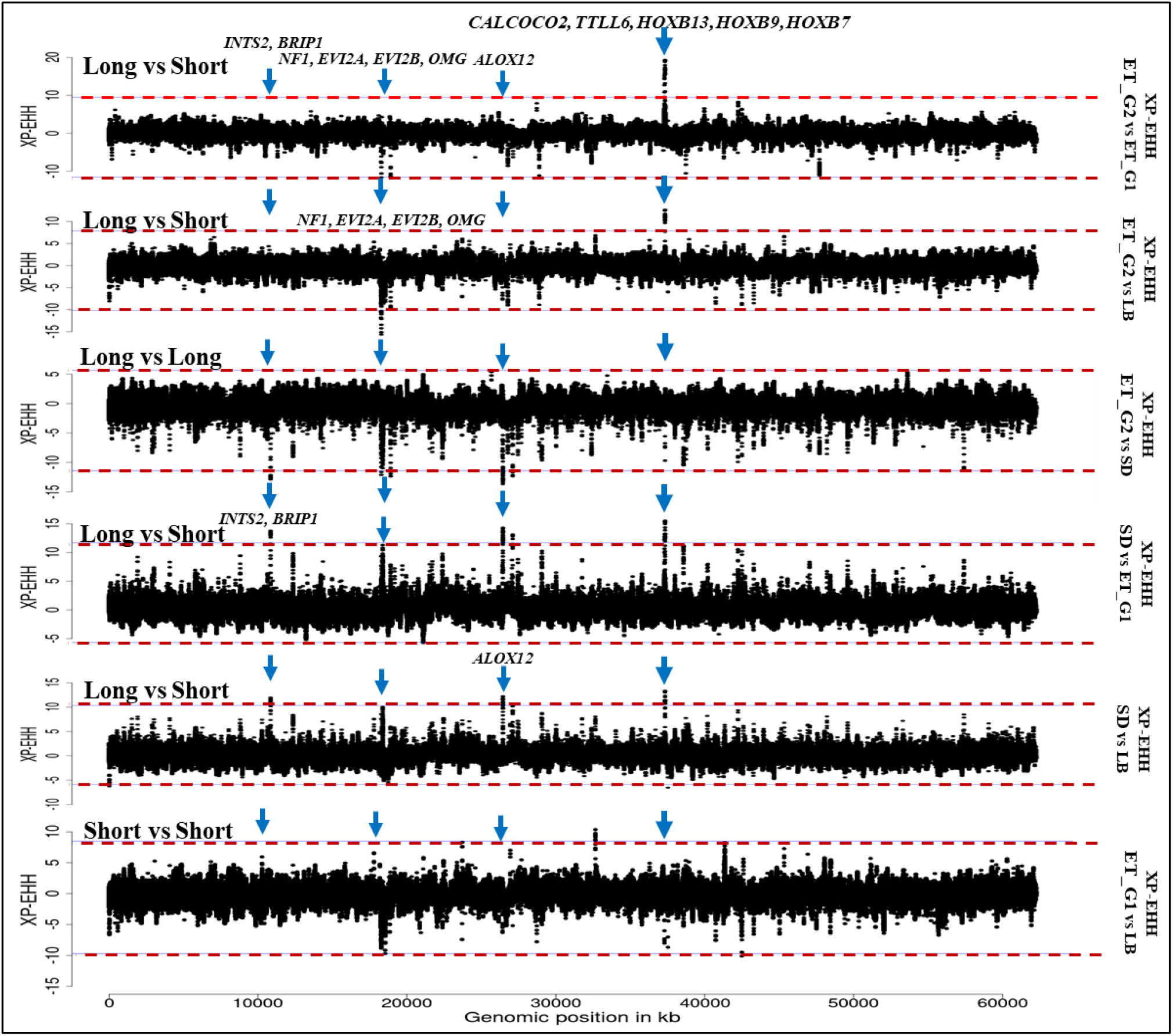
Signatures of selection revealed by XP-EHH on OAR11 spanning *HOXB13*, previously identified in connection with tail length as well as new candidate regions spanning genes related to fat metabolism (e.g., *ALOX12, NF1, EVI2B* and *OMG*).

**Figure 11.**
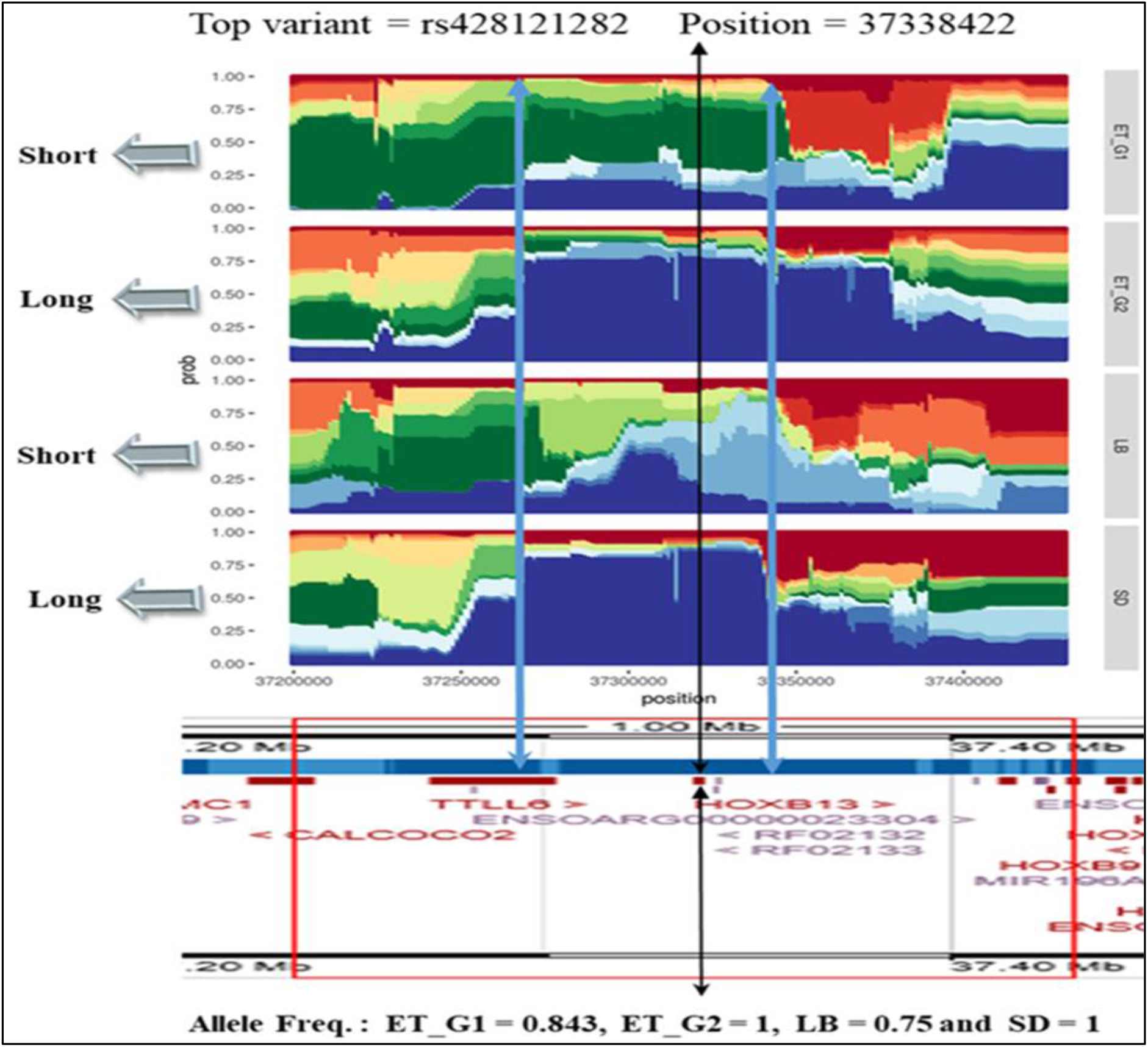
The haplotype structure around the candidate region on OAR11 spanning *HOXB13* likely associated with vertebrae length.

To identify selection signatures for tail fat-depot sizes, we contrasted the thin-tailed (SD) sheep with the fat-rumped (ET-G1) and the two fat-tailed groups (LB and ET-G2). The three approaches (*Hp*, *FST*, *XP- EHH*) identified 34 candidate regions overlapping 122 genes. Twelve regions were identified by one of the three approaches and 25 by a combination of at least two approaches (Table S13; Figure 12). Table 3 shows the candidate genes associated with fat deposition and which were present in the candidate regions identified by our analysis.

**Figure 12.**
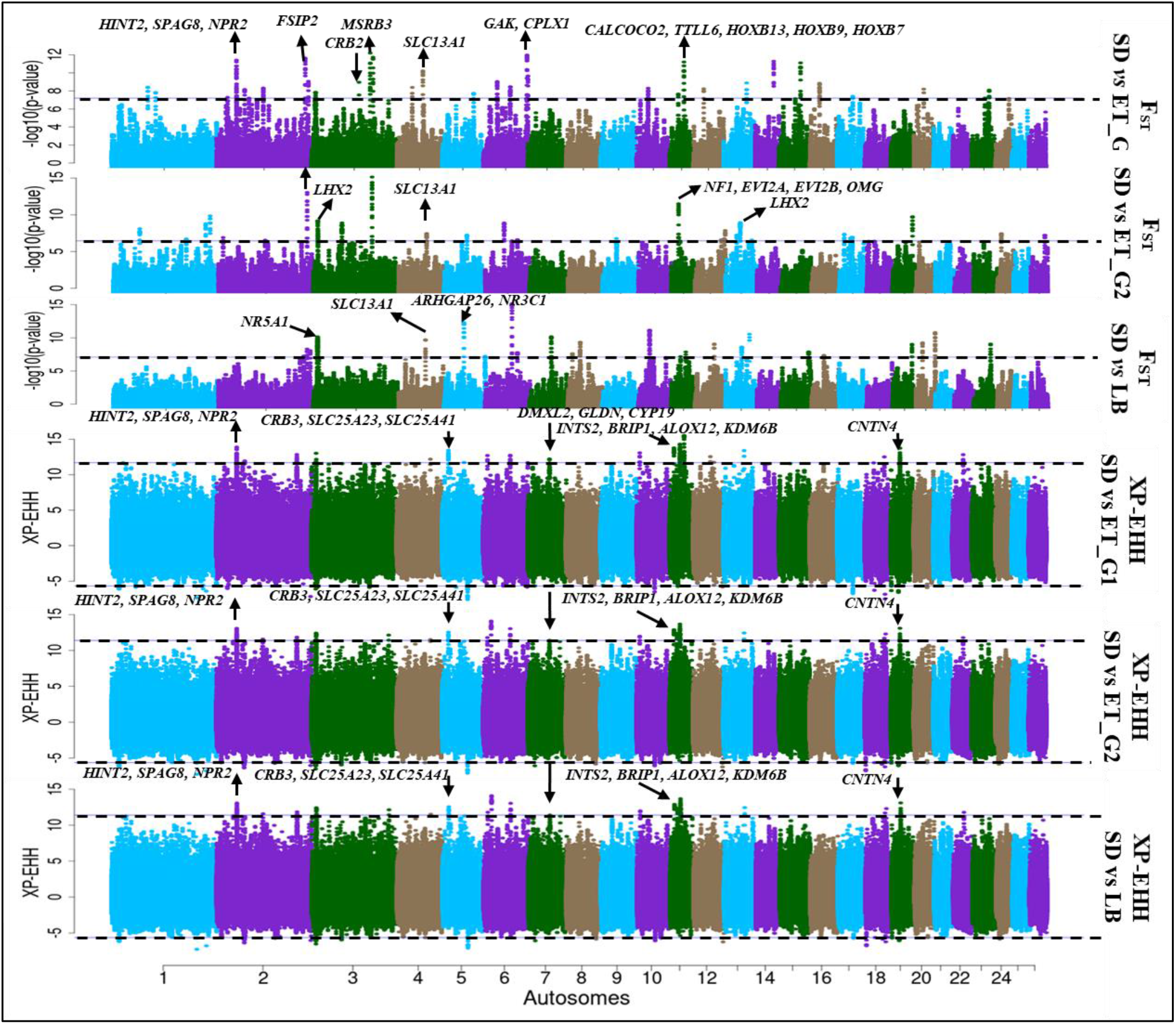
Manhattan plot showing genome-wide distribution of FST and XP-EHH values resulting from contrasting of thin-tail SD sheep group with fat-rumped (ET-G1) and fat-tailed (ET-G2 and LB) sheep groups.

## Discussion

### Population structure and demographic history

Human and natural selection have shaped the genomes of domestic livestock since domestication. Here we investigated the genome architecture of indigenous African sheep, by analysing whole-genome sequences of 150 indigenous sheep from three African countries, Ethiopia, Sudan and Libya. We compared the sequence reads to the OAR_v3.1.75 reference genome assembly and obtained 34.85 million high quality variants. An average of 93.62% of the SNPs were validated in the sheep dbSNP database of which 0.79% were exonic while the majority (55.9%) were intergenic. This is consistent with observations in Eurasian wild aurochs (Park et al., 2015) and Chinese native sheep (Yang et al., 2016), indicating high quality of our sequences, high reliability of the called SNPs and completeness of the gene regions.

Although the history of indigenous African sheep has been inferred from archeological studies, the picture still remains incomplete. Our phylogenetic inference identified four distinct genetic groups underlying the genetic structure of our study populations. These genetic groups are offering interesting insights on the historical diffusion and dispersal of sheep into and across the African continent. The North African Libyan Barbary (LB) and the North-East African Sudanese Hammari and Kabashi (SD) sheep clustered in close proximity but separate from each other and the East African sheep represented here by Ethiopian sheep; the latter separated into two genetic groups, ET-G1 and ET-G2. This result supports for the first time, based on whole genome sequence data, two distinct entry points of sheep in the continent, the Isthmus of Suez (LB and SD) and the Horn of Africa (ET-G1 and ET-G2). The sub- grouping of Ethiopian sheep into ET-G1 (fat-rumped) and ET-G2 (long fat-tailed) is indicative of different expansion events through the Horn of Africa. Based on the current dataset, it is difficult to infer whether ET-G1 and ET-G2 arrived together or independently. However, the geographic distribution of the Ethiopian populations in each group offers indirect insights. ET-G1 populations are found in areas close to the Red Sea and Indian Ocean coastline, while ET-G2 ones occur in the country’s hinterland. The ancestral populations of these two groups may therefore have arrived independently, with the arrival of the long fat-tailed sheep preceding those of the fat-rumped.

Three ancestral backgrounds were supported following ADMIXTURE analysis. All three are present in Sudanese sheep which also show the highest *N*e (∼2720) 1000 generations ago. The ancestor of domestic sheep was thin-tailed, but its tail was shorter than that of the domestic variant. Accordingly, as supported by archaeological thin-tailed iconography on the African continent, the thin-tailed sheep are likely to be the most ancient of the African sheep lineages (Muigai and Hanotte, 2013). The Sudanese and Libyan sheep shared a large proportion of their genome ancestry and show close genetic proximity on the PCA. They are likely the modern-day legacy of the ancient arrival of thin-tailed sheep on the African continent followed by their dispersion along the Mediterranean Sea coast (Libya) and southwards along the Nile River Basin (Sudan). Although the ET-G2 and LB sheep are both fat-tailed, they do have different ancestral backgrounds, supporting two entry points for African fat-tailed sheep on the continent i.e., the Isthmus of Suez and the Horn of Africa.

The ADMIXTURE analysis also revealed that three populations, Doyogena, Shubi Gemo and Gafera, of ET-G2 share ∼15-20% of their genome ancestry with ET-G1. Likewise, we detect the main ancestry of ET-G2 in three populations of ET-G1 (Menz, Adane, Arabo). This pattern of intermixing can be attributed to flock movements/exchanges arising from human socio-cultural and economic activities (Gizaw et al., 2007). This is supported by TreeMix which provided evidence of gene flow from fat- rumped sheep from eastern Ethiopia into long fat-tailed sheep in central and southern Ethiopia. A similar pattern of admixture was observed between six Ethiopian populations (Menz, Gafera, Gesses, Kido, Adane and Arabo) and the Sudanese (SD) sheep. This admixture pattern is consistent with previous findings (Ahbara et al., 2019). Half of the individuals of LB shared ∼30% of their genome ancestry with those of SD and ET-G1. We suggest this shared ancestry is the result of inter-crossing fat-tailed with thin-tailed sheep to reduce the volume of tail fat in the former due to the decline in the monetary value of animal fat (Moradi et al., 2012; Moioli et al., 2015). Such inter-crossings have been reported in Algeria (Gaouar et al., 2017) and Tunisia (Rekik et al., 2005). It is worth noting that large fat tails, which are discarded following slaughter, can account for up to 14.5% of the cold dressed carcass weight in sheep (Kiyanzad, 2005). The shared ancestries between the north African SD and LB sheep in line with the equal number of tail vertebrae (Figure 1a) and different tail sizes may support such pattern of intermixing. On the other hand, various ancestries and varying numbers of tail vertebrae paired with diverse tail sizes might imply a different history of Ethiopian sheep (Figure S4b).

### Genome targets for adaptation to environments

The African landscape has pronounced changes in elevation ranging from sea level and rising to high altitudes in the East African highlands. The variation in altitude is accompanied by large variations in agro-climatic conditions ranging from dry and arid to wet and humid. Such variations in topography and environmental conditions allow studying adaptive processes in response to environmental selection pressure, e.g., hypobaric hypoxia in high altitudes, and heat, water and feed stress in the drylands. It provides excellent opportunities to investigate how natural selection shapes the genetic architecture of adaptive divergence. Here, we sampled 15 populations of indigenous African sheep from across the continents ecological and altitudinal range. We observed a separation of the populations into four genetic groups that align well with geography. Furthermore, our comparative genomic analyses involving the four genetic groups revealed several important observations shedding new insights on their biology of adaptation.

Hypobaric hypoxia imposes major stresses on the physiology of organisms at high altitudes and a powerful homeostasis system has evolved to offset ambient hypoxia and attain physiological homeostasis (Rytkönen and Storz, 2011). The classic response is an increase in haemoglobin concentration to compensate for the unavoidable lowered percent of oxygen saturation. Our selective sweep mapping involving ET-G2 identified candidate regions spanning nine genes, *TF* (OAR1), *EPAS1* and *ADSL* (OAR3), *NF1* (OAR11), *PLCG1* and *CHRNA4* (OAR13), *FGF2* (OAR17), *VEGFA* (OAR20) and *EGLN1* (OAR25). These genes comprised the two most significant GO terms, “response to hypoxia” and “positive regulation of angiogenesis”, and KEGG pathways, “HIF-1 signalling pathway” and “pathways in cancer”, suggesting they may underlie adaptive changes in high-altitude environments. Indeed, *EGLN1* and *EPAS1* encode the synthesis of prolyl hydroxylase domain 2 (*PHD2*) that plays a critical role in adapting to changes in oxygen levels and its variation has consistently been associated with haemoglobin levels in Tibetans (Simonson et al., 2010). It is also a target of natural selection for hypoxia adaptation in humans and animals (Storz and Moriyama, 2008; Tissot Van Patot and Gassmann, 2011). The HIF-1 pathway plays a central role in regulating cellular responses to hypoxia (Frede and Fandrey, 2014) in several species found at high altitudes (see review by Friedrich and Wiener (2020)). The *NF1* gene occurred in a candidate region under selection in sheep living at high altitudes (Wei et al., 2016; Yang et al., 2016). *NF1* promotes and regulates angiogenesis in response to hypoxia on its own (Wu et al., 2006) or as a component of the Ras/ERK and VEGF signalling pathways (Minet et al., 2000). *VEGFA* has been associated with cerebrovascular adaptation to chronic hypoxia in sheep (Castillo-Melendez et al., 2015) and protects pregnant ewes from hypoxic effects by increasing blood flow in uterine arteries (Mehta et al., 2012). Under hypoxic conditions, elevated expression of *TF* and *VEGFA* increases oxygen delivery by stimulating iron metabolism and angiogenesis, following a positive feedback from *EGLN1*. Unexpectedly, our analytical strategy to identify genomic regions associated with differences in tail length and tail fat-depot size also identified selection targets relating to high altitude adaptation. For instance, the gene encoding methionine sulfoxide reductase B3 (*MSRB3*) occurred on a candidate region on OAR3 that was supported by *FST* and *XP-EHH*. This gene has been associated with high altitude adaptation in dogs (Gou et al., 2014) and Tibetan sheep (Wei et al., 2016). Our results appear to suggest that, driven by a hypoxic environment in the Ethiopian highlands, natural selection has been acting on a group of genes associated with the HIF-pathway and angiogenesis in driving adaptive processes in the ET-G2 group. Although we do find compelling evidence implicating a genetic predisposition to high altitude adaptation in ET-G2, previous studies in humans (Alkorta-Aranburu et al., 2012), cattle (Edea et al. 2014) and sheep (Edea et al., 2019) in Ethiopia did not find similar associations with the HIF-pathway and angiogenesis. These contrasting findings are rather surprising, but we speculate that they could be the result of differences in sampling strategies, analytical approaches and the markers used. For instance, the studies by Edea et al. (2014; 2019) and Wiener et al. (2021) classified their study populations based on geographic sampling; Wiener et al, also used an ecological approach to analyse their data. In contrast, we used the genetic groups revealed by the phylogenetic inference to inform our analysis.

Haplotype analysis around the candidate regions associated with high altitude adaptation returned some unexpected results. The haplotype structures around the *TF* locus showed a homogeneous haplotype approaching fixation in ET-G2, but several segregating haplotypes in SD and LB (Figure 4b), suggesting selection around the locus in ET-G2. Haplotype structure analysis around *EGLN1* showed SD shared identical haplotypes with ET-G2 that appeared to be approaching fixation, against an admixed haplotype in LB. However, LB and ET-G2 shared a similar selection sweep occurring around the *NF1* gene (Table 2; Figure 3,5). In Egyptian sheep that are adapted to a dryland environment, Mwacharo et al. (2017) found candidate regions spanning genes that were highly enriched for the GO term “response to hypoxia” and “HIF-1 signaling pathway”. They suggested this could be an adaptive response to physical exhaustion due to long-term long-distance trekking in search of pasture and water resources, an activity that results in hypoxia-like conditions and oxygen debt in skeletal muscles. These results suggest that parallel evolution driven by distinct natural selection forces could be acting on the same set of genes driving adaptation to different environments in the same species. This is rather surprising and calls for further investigations.

High ambient temperatures resulting in thermal stress, present the physiological challenge to organisms residing in hot arid environments. Thermal stress impacts production and reproduction and desert dwelling animals have to invoke physiological responses to mitigate thermal stress and its negative effects. The analysis involving SD and LB sheep groups identified 123 candidate regions spanning 182 genes (Table S10); 19 were reported in candidate regions revealed in Chinese sheep living in desert and arid environments (Yang et al., 2016). Among the genes identified in a candidate region on OAR11 was *KCNAB3* which encodes a potassium voltage-gated channel subunit that is correlated with cognitive performance under chronic stress (Jung et al., 2017) and is down-regulated in response to acute and chronic stress in mice (Terenina et al., 2019). This result is enlightening given that LB and SD sheep are exposed to extended periods of thermal stress in Libya and Sudan, respectively. Apart from thermal stress, high temperatures constrain pasture availability both in quality and quantity. One of the candidate regions in which SD and LB shared a similar haplotype, occurred in OAR2 and spanned *DIS3L2* (Figure 6a). This gene was found to be a candidate under selection in cattle (Gautier et al., 2009) and Brazilian sheep and has been associated with height variation (de Simoni Gouveia et al., 2017). Given that large animals generally have higher maintenance requirements (Gomes et al., 2013), the occurrence of *DIS3L2* in a candidate region in LB and SD may act to regulate their physical size while providing the advantage of better thermoregulation (McManus et al., 2011).

Thermal stress affects male reproductive performance through decreased semen volume, sperm motility, higher sperm defects and reduced libido (see review by Sejian et al. (2019). We found *CNTROB* (centrobin, for centrosome BRCA2 interacting protein) in a candidate region on OAR11. Through its role in spermiogenesis (Pandey et al., 2019), *CNTROB* is critical in maintaining male fertility (Liška et al., 2013). One region that differentiated LB from ET-G2 and SD and showed a distinct haplotype approaching fixation in LB (Figure 6d) presented on OAR10 and spanned *PDS5B* (PDS5 cohesin associated factor B). This gene modulates cohensin functions in spermatocytes and spermatogonia, contributing to meiotic chromosome function and structure (Fukuda and Hoog, 2010). By virtue of occurring in a selective sweep region in dryland dwelling sheep, *CNTROB* and *PDS5B* may be critical in maintaining male fertility and thus reproductive success under prolonged heat stress.

Soil salinisation contributes to soil degradation and compromises water and pasture quality (McFarlane et al., 2017). Droughts, which exacerbate soil salinity, have increased over the years due to global warming (Busby et al., 2010; Bindra et al., 2013) and worsened in arid and semi-arid environments (McFarlane et al., 2017). Adaptive responses to increased salinity and droughts in drylands may explain the selection signatures spanning *PLEKHA7* (Pleckstrin homology domain containing A7) in a region on OAR15 that differentiates LB (Figure 6c). *PLEKHA7* is highly expressed in kidney and heart (Pulimeno et al., 2010); it was reported to attenuate salt-sensitive hypertension and renal diseases in rats (Endres et al., 2014). In humans, *PLEKHA7* has been associated with blood pressure, kidney function and hypertension (Lin et al., 2011; Taal et al., 2012). Two of the most significant KEGG pathways were “Aldosterone synthesis and secretion (GO:0030325)” and “Vasopressin-regulated water reabsorption (oas04962)”. Aldosterone and vasopressin regulate blood pressure and maintain electrolyte and water balance thus facilitating homeostatic adaptations (Harvey et al., 1984). The overexpression of *NF1*, which we found in one of the candidate regions under selection, results in reduced transcription of *HSD11B2* possibly associated with salt sensitivity (Alikhani-Koupaei et al., 2007) and hypertension (Friedman et al., 2002) in humans. *NR5A1* plays a critical role in the expression of *CYP11B2* (Takeda et al., 2018) which plays a key role in salt-sensitive hypertension development (Fujita, 2008) and in aldosterone biosynthesis in the adrenal cortex (Curnow et al., 1991) in humans.

### Selection targets for phenotypic traits

We observed a candidate region on OAR11 that was specific to SD and ET-G2 groups. This region included several genes among which four, *TTLL6, HOXB13, HOXB9* and *HOXB7* have been associated with growth and development of the spinal cord and tail vertebrae in several animal species (Burke and Nowicki, 2001; Economides et al., 2003; Mallo et al., 2010; Ahbara et al., 2019). This region occurred upstream of two candidate regions spanning five genes (*NF1, EVI2A, EVI2B, OMG* and *ALOX12*) (Figure 10, 12) that have been associated with fat deposition (Moioli et al., 2015; Wei et al., 2015; Yuan et al., 2017; Ahbara et al., 2019). The fact that we observed two regions or more occurring adjacent to one another and associated with two tail traits (fat-depot size and CV length) suggests a natural selection phenomenon influencing tail traits in sheep. One region on OAR3 spanned *NR5A1* and *NR6A1* which influence vertebrae number in pigs (Mikawa et al., 2007; Rubin et al., 2012) and thoracic vertebrae number in sheep (Shengwei et al., 2019). A candidate region spanning *PDGFRA* (platelet derived growth factor receptor alpha) was also identified by the three methods on OAR6 in LB and a distinct haplotype was found around this gene. *PDGFRA* is involved in the differentiation of preadipocytes and was also found in a candidate selection region associated with the fat-tailed phenotype in sheep (Mastrangelo et al., 2019). These results suggest possible co-evolution between tail skeleton morphology and tail fat deposition in sheep.

Unexpectedly, our analytical strategy that aimed to identify candidate regions associated with differences in tail length and tail fat-depot sizes, identified selection targets relating to other traits. For instance, the gene encoding methionine sulfoxide reductase B3 (*MSRB3*) occurred on a candidate region on OAR3 supported by *FST* and *XP-EHH*. An association between *MSRB3* with ear types has been reported in dogs (Boyko et al., 2010; Vaysse et al., 2011), sheep (Wei et al., 2015) and pigs (Chen et al., 2018). An examination of haplotype structures around *MSRB3* showed a pattern that was consistent with the ear phenotypes of the three sheep groups analysed here. Compared to individuals of LB and SD, majority of which have long drooping ears, the ET-G2 represented by Doyogena, revealed a large, conserved haplotype around *MSRB3*. This result is consistent with the observation that polytypic species usually exhibit shorter extremities such as limbs and ears in cold environments while the ones inhabiting warmer climates show longer extremities (Allen, 1907), suggesting such differences are indicative of environmental adaptation.

LB sheep differentiated from ET-G2 at two candidate regions on OAR10. One of these spanned *RXFP2* (relaxin family peptide receptor 2) which determines horn size and development in ungulates (Kardos et al., 2015) and primary sex characters in humans and mice (Johnston et al., 2011). Heat-exchange capacity shows an evolutionary trend of variation in horn core size in response to ambient temperatures (Hoefs, 2000). Although the selective sweep spanning *RXFP2* could implicate its dual role in the development of horns for thermoregulation and enhancing reproductive success under thermal stress, in recent times sheep farmers in Libya are preferring breeding rams with larger horns and tail sizes because such rams are believed to result in strong and healthier progenies (Figure S6). The selection signature around *RXFP2* may therefore be driven by human selection for horn size and natural selection for the thermoregulation function of horns, although the semi-feral nature of management of LB sheep could also be implicated as suggested for Chinese sheep (Pan et al., 2018).

### Selection targets for resistance to endo- and ecto-parasites

High altitude hypoxia increases vulnerability to infections by decreasing the secretion of cytokines and immunoglobulins (Mishra and Ganju, 2010). In arid environments, feed, water and heat stress can also weaken immunity (Silanikove 2000). The candidate region on OAR11 that spanned four genes (*NF1*, *EVI2A*, *EVI2B*, *OMG*) associated with tail morphology, has also been linked to gastrointestinal nematode infection resistance in traditionally managed Tunisian sheep (Ahbara et al., 2021). The *PDGFRA* gene that occurred in a candidate region on OAR6, and which has been associated with the fat-tail phenotype (Mastrangelo et al., 2019) likely through the differentiation of preadipocytes, has also been implicated in host immune responses against endoparasites and is a key gene in cytokine signaling (Beh et al., 2002; Benavides et al., 2015). The *Hp* analysis detected a candidate region on OAR1 and OAR6 in ET-G1, ET-G2 and SD sheep. The region on OAR1 includes *CD84*, reported as a candidate gene for trypanosomiasis resistance in mice (Yan et al., 2004; Martin and Tarleton, 2005). The region on OAR6 includes *LETM1* which plays an essential role in the infection stages of *Trypanosoma brucei* by maintaining its mitochondrial integrity (Nowikovsky et al., 2012; Hashimi et al., 2013; Nowikovsky and Bernardi, 2014). These regions appear to be maintained in ET-G1, ET-G2 and SD tropical sheep which may have been exposed to endo-parasite and trypanosomosis challenge at the opposite of the North African LB sheep (Table 4). Further investigations are required to confirm the significance of these results in relation to resistance/tolerance to nematode infections and trypanosomiasis.

**Table 4.**
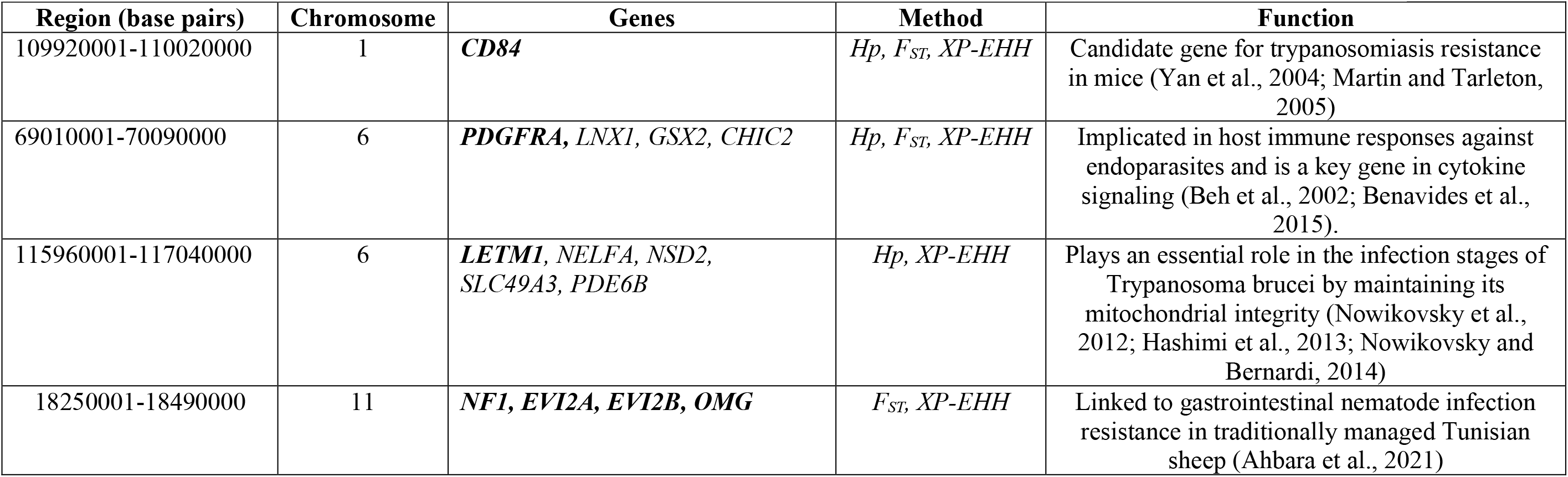
Candidate regions and genes associated with resistance to endo- and ecto-parasites as detected in our study.

### Possible pleitropic effects

Taken together, it may be argued based on our results that the interaction between genes, whether on the same or different chromosomes, driven by environmental conditions are responsible for adaptation to diverse environments and the diversity in sheep tail phenotypes. For example, on OAR11 the occurrence of *KDM6B* within a candidate region that is close to other regions associated with fat accumulation may regulate the expression of *HOXB13* (Figure 10).

## Conclusion

Our results revealed four autosomal genomic backgrounds in indigenous East, North and North-East African sheep. The genotypes of most of the individuals analyzed here comprised at least two genetic backgrounds which could be accounted for by some level of current and/or historical intermixing. Our findings corroborate archaeological evidence that have revealed domestic sheep migration events and entry points into Africa. Besides, our results support possible interaction between environmental factors and tail morphology in shaping the genomes of indigenous sheep inhabiting diverse environments. From the perspective of environmental adaptation, our study populations illustrated the genome structure towards environmental challenges (coastal hot-humid arid, hot-dry desert and high-altitude humid). Our whole-genome sequence analysis also suggest interaction between several genomic regions for tail phenotype in sheep. For instance, many candidate genes possibly under selection were identified, among which, the salt-sensitivity related gene (*PLEKHA7*) in Libyan sheep, thermoregulation genes (*CREB3L2, CREB3, GNAQ, DCTN4*) in Libyan and Sudanese sheep, hypoxia associated genes (*EGLN1*, *EPAS1*) in Ethiopian sheep, and skeletal tail associated genes (*HOXB13, NR5A1*, *NR6A1, KDM6B* and *NR3C1*) most likely are playing major roles in the unique adaptations of African sheep. To the best of our knowledge, we are the first to report these candidate genes in African indigenous sheep.

However, future studies including thin-tailed sheep from Ethiopia and both thin- and/or fat-tailed sheep from West, northern and southern Africa could provide a comprehensive view on the genome architecture of African indigenous sheep.

## Data Accessibility

The datasets analyzed in this study are available upon request from the corresponding authors.

## Ethics statement

The animals used in this study are owned by farmers. Prior to sampling, the objectives of the study were explained to them in their local languages for them to make informed decisions regarding giving consent to sample their animals. Government veterinary, animal welfare and health regulations in Ethiopia, Libya and Sudan were observed during sampling. Skin tissues importation and/or exportation was permitted by the Ethiopian Ministry of Livestock and Fisheries under Certificate No: 14-160-401-16.

## Author Contributions

AMA, OH, and JMM conceived and designed the study. AMA analyzed the data with inputs from JMM and wrote the manuscript. OH and JMM revised the manuscript. EC, CR, and PW contributed to sequencing and mapping data of Ethiopian and Libyan sheep breeds and provided critical inputs on data analysis. HM contributed to sequencing samples of Sudanese sheep. AA, SL and MOA contributed and coordinated the sampling in Ethiopia and Libya, respectively. All authors read and approved the final manuscript.

## Acknowledgments

This study was conducted during AMA’s PhD training which was supported by the Libyan Ministry of Higher Education and Scientific Research and the University of Misurata. Sampling of Ethiopian sheep was supported by the CGIAR Research Program on Livestock. Accordingly, ICARDA and ILRI wish to thank the donors and organizations that globally support the work of the CGIAR Research Program on Livestock through their contributions to the CGIAR Trust Fund. Sampling of Libyan sheep was supported by the Libyan Agriculture Research Centre (Misurata station) and the University of Misurata (Department of Animal Production).

---

**Figure S1.**
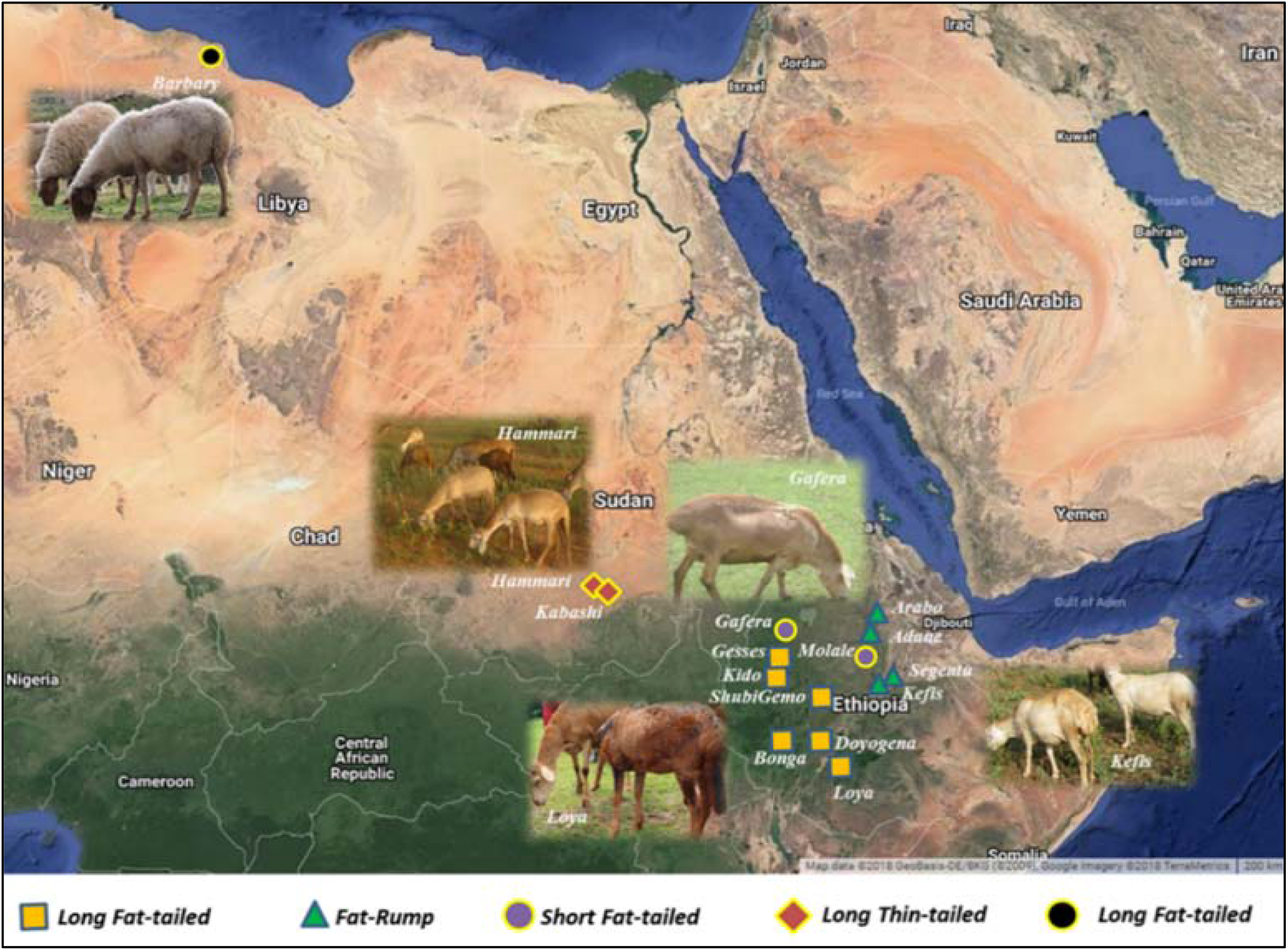
The sampling locations of Ethiopian, Sudanese and Libyan sheep populations.

**Figure S2.**
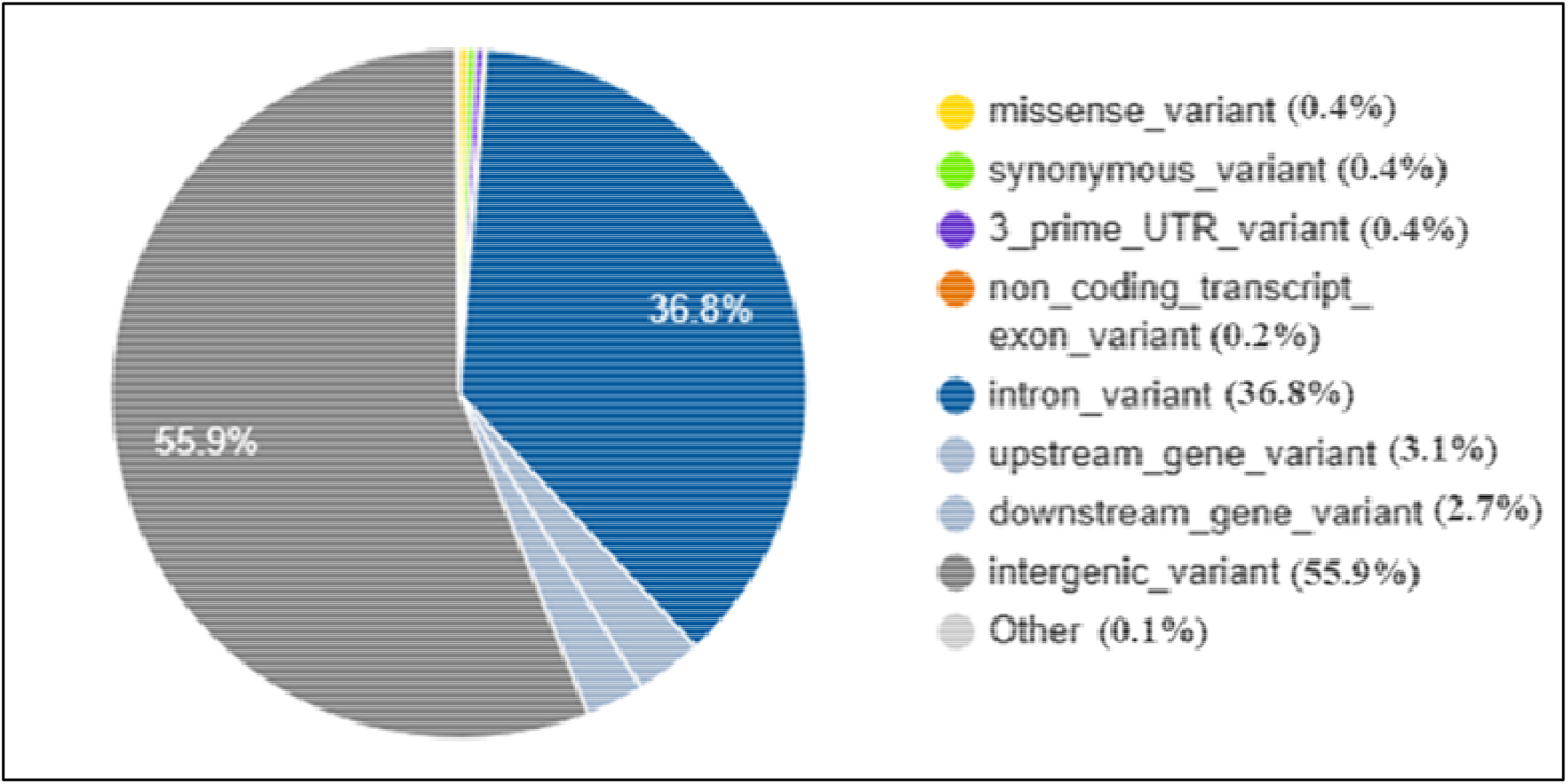
Distribution of SNPs following annotation categories (150 sheep samples).

**Figure S3.**
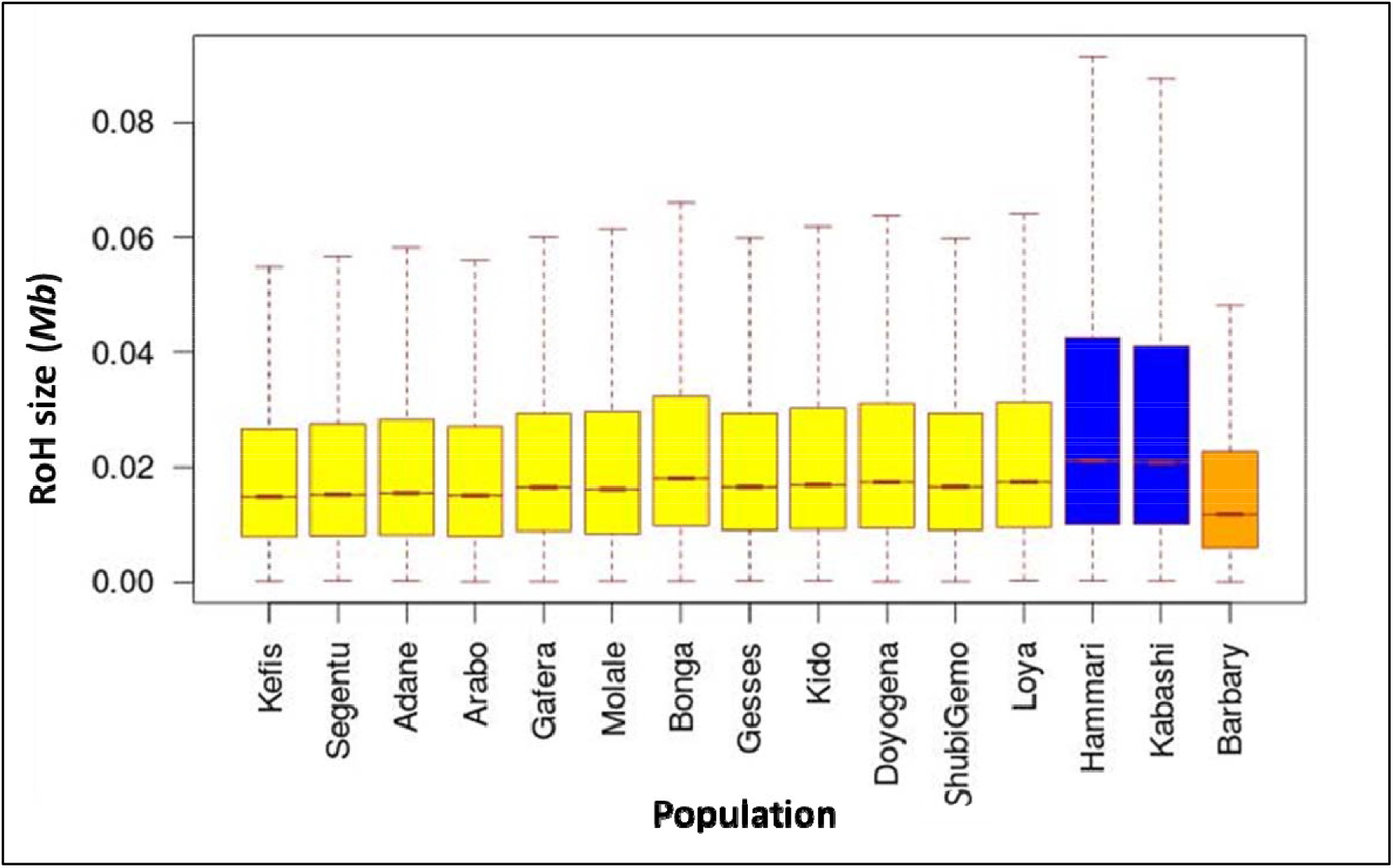
Distribution of RoHs within range of 1.4 Mb for each sheep population.

**Figure S4.**
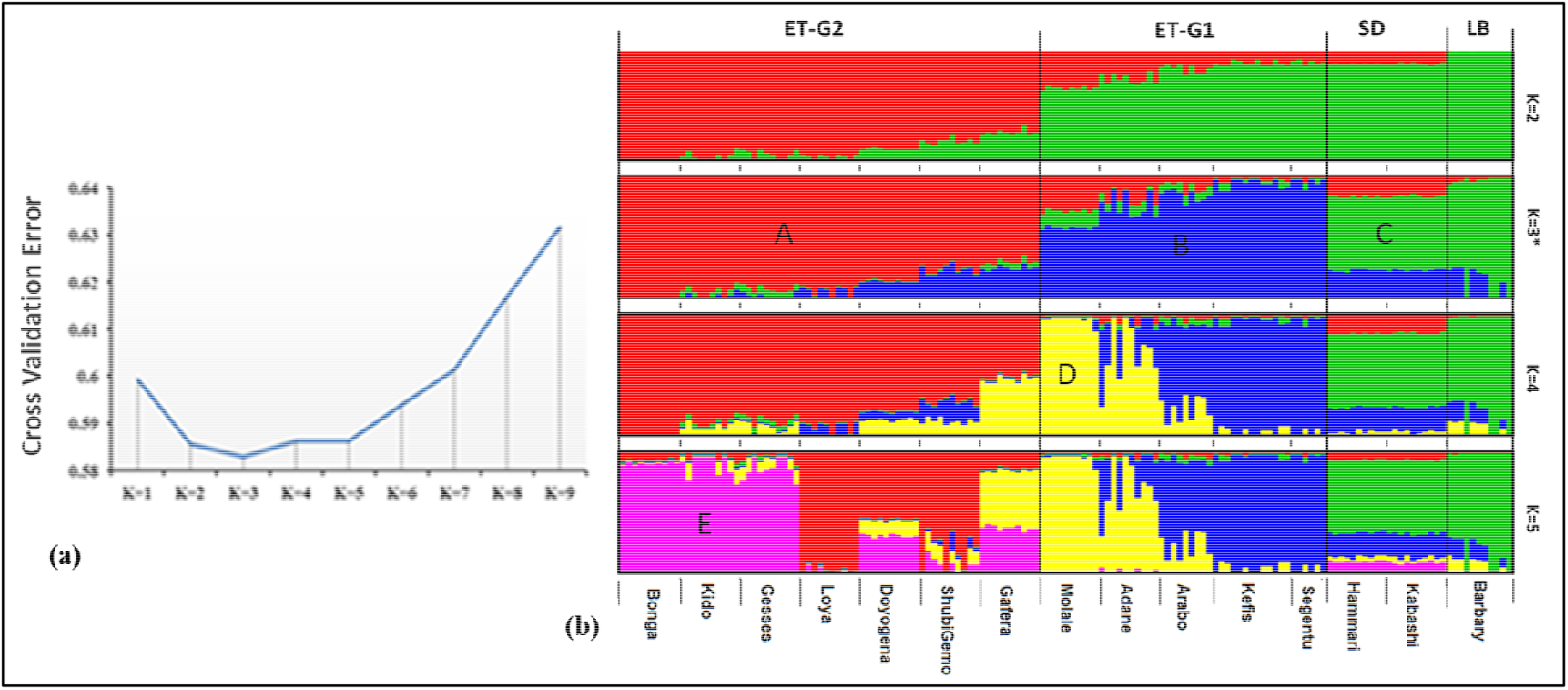
Genetic admixture analysis of the studied populations (a) Cross-validation plot (b) Admixture plot.

**Figure S5.**
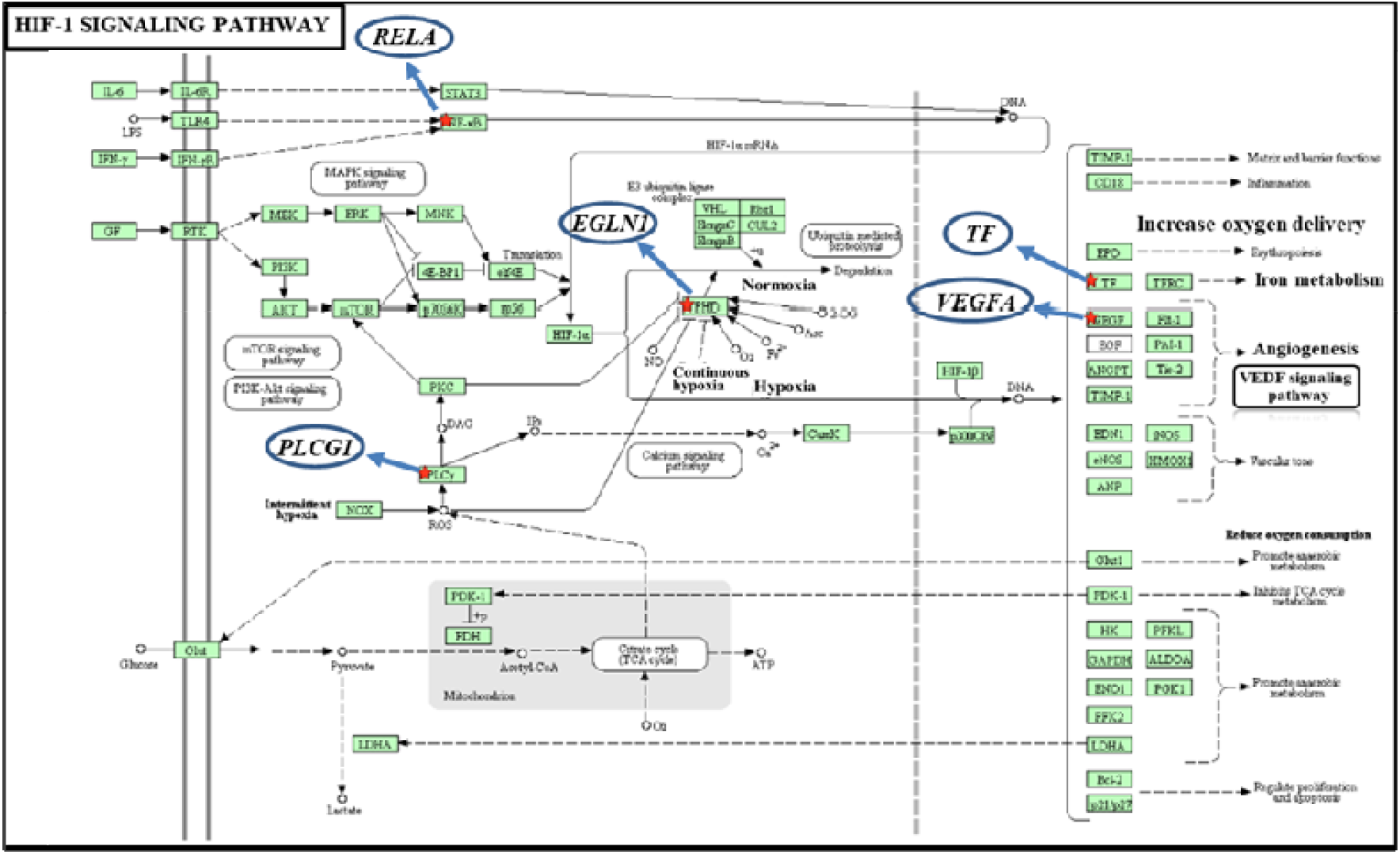
Plot of HIF-1 pathway showing the key role of the positively selected genes for high-altitude adaptation in African sheep.

**Figure S6.**
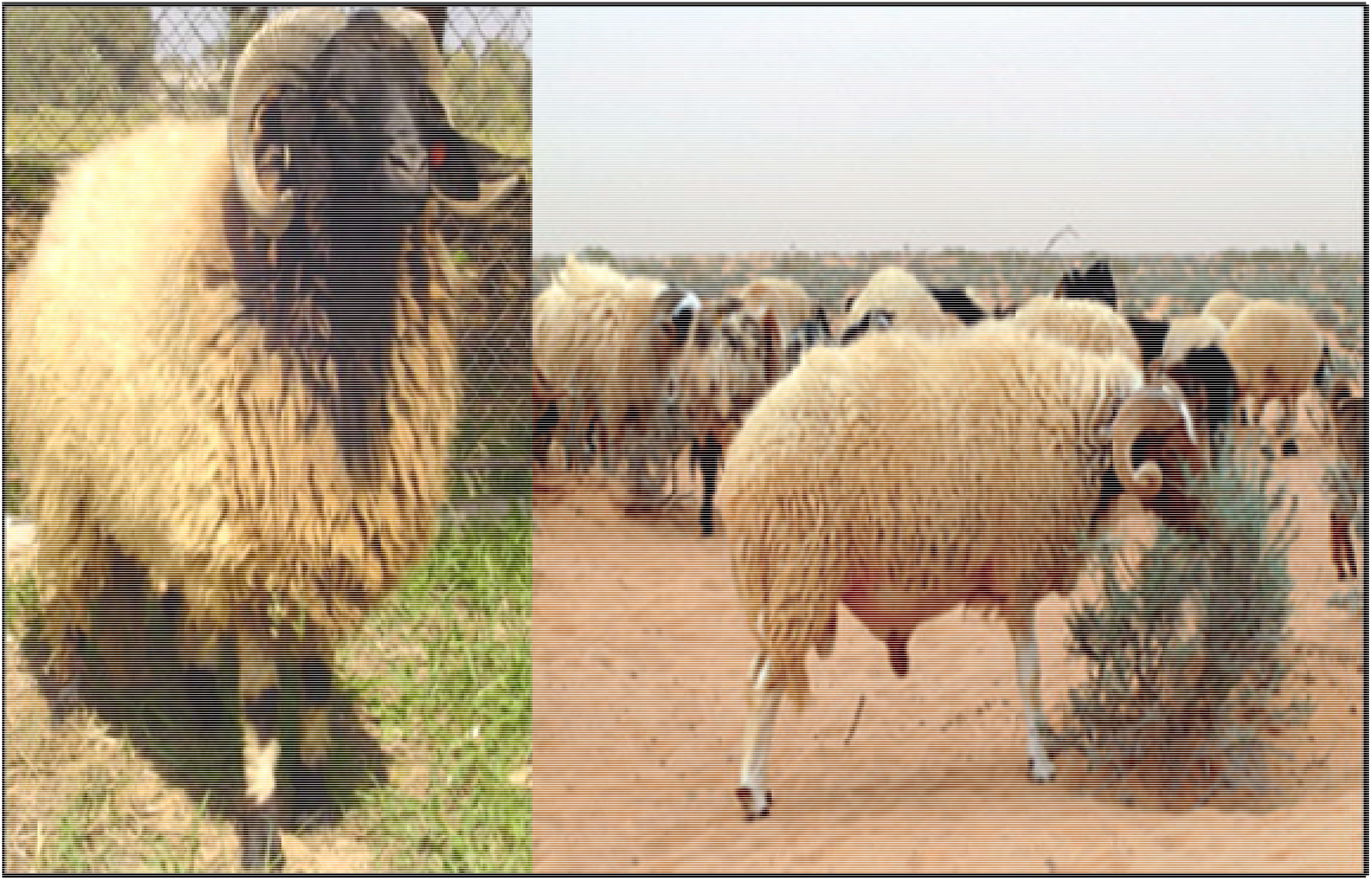
Illustration of the phenotypic criteria (horns and tail) applied by the owners of sheep in Libya when selecting ram for reproduction.

