## Supplementary Tables for "The natural adaptation and human selection history of African sheep genomes"

**Table S1** Geographic origin and tail types of the studied populations

| **Origin** | **Population** | **Latitude (*N*)** | **Longitude (*E*)** | **Altitude (meter asl)** | **Sample size** | **Tail Type** |
| --- | --- | --- | --- | --- | --- | --- |
| **Ethiopia** | Kefis | 9°30’ | 40°10’ | 890 | 13 | Fat-Rumped |
|  | Segentu | 9°27’ | 40°05’ | 740 | 6 | Fat-Rumped |
|  | Adane | 11°14’ | 39°50’ | 2450 | 10 | Fat-Rumped |
|  | Arabo | 11°09’ | 39°54’ | 1500 | 9 | Fat-Rumped |
|  | Gafera-Washera | 11°31’ | 36°54’ | 2500 | 10 | Short fat-tailed |
|  | Molale/Menz | 10°70’ | 39°39’ | 3068 | 10 | Short fat-tailed |
|  | Bonga | 7°16’ | 36°15’ | 1788 | 10 | Long fat-tailed |
|  | Gesses | 10°50’ | 36°14’ | 1300 | 10 | Long fat-tailed |
|  | Kido | 10°71’ | 36°19’ | 1300 | 10 | Long fat-tailed |
|  | Doyogena | 7°21’ | 37°47’ | 2324 | 10 | Long fat-tailed |
|  | Shubi Gemo | 8°80’ | 38°51 | 1600 | 10 | Long fat-tailed |
|  | Loya | 6°29’ | 38°24’ | 1900 | 10 | Long fat-tailed |
| **Sudan** | Hammari | 13°09’ | 29°22’ | 620 | 10 | Long thin-tailed |
|  | Kabashi | 13°09’ | 29°22’ | 620 | 10 | Long thin-tailed |
| **Libya** | Barbary | 32°31’ | 15°09’ | < 100 | 12 | Long fat-tailed |
|  | | | | | **150** |  |

**Table S2** Classification of sheep groups and populations in contrasting environments based on their tail type, genomic group, altitude and environment categories.

| **Origin** | **Tail type** | **Genomic group** | **N** | **Altitude category** | **Range (meter asl)** | **Environment category** | **Precipitation (mm)** | **Populations** |
| --- | --- | --- | --- | --- | --- | --- | --- | --- |
| Libya | Long fat-tailed | LB | 12 | Sea level | < 50 | Hot semi-arid | 252.6 | Libyan Barbary |
| Sudan | Long thin-tailed | SD | 20 | Low altitude | < 620 | Hot arid and desert | 233.3 | Hammari, Kabashi |
| Ethiopia | Long fat-tailed | ET-G2 | 70 | High altitude | 1300 - 2500 | Humid and sub-humid | 1144.1 - 1787 | Gesses, Kido, ShubiGemo, Bonga, Loya, Doyogena, Gafera (short fat-tailed) |
|  | Fat-rumped | ET-G1 | 19 | Low altitude | 740 - 890 | Hot arid lowland plain | 404.5 | Segentu, Kefis |
|  |  |  | 29 | High altitude | 1500 - 3068 | Cool to very cold sub-moist/dry alpine mountains and plateaus | 1102 | Arabo, Adane, Molale (short fat-tailed) |

**Table S3** SNP statistics for each sheep population

| **Population** | **Total number of SNPs** | **Average per sample** | **Het/Hom** | **dbSNP (%)** |
| --- | --- | --- | --- | --- |
| Kefis | 37,531,090 | 12,561,119 | 2.15 | 93.7 |
| Segentu | 25,436,133 | 12,472,311 | 1.65 | 93.5 |
| Adane | 26,713,625 | 12,574,438 | 2.13 | 93.5 |
| Arabo | 28,466,085 | 12,617,954 | 2.03 | 93.5 |
| Gafera-Washera | 34,128,747 | 12,398,645 | 1.98 | 93.7 |
| Molale-Menz | 34,547,334 | 12,401,271 | 2.10 | 93.7 |
| Bonga | 34,274,566 | 12,152,710 | 1.96 | 93.7 |
| Gesses | 34,076,190 | 12,400,542 | 1.87 | 93.7 |
| Kido | 33,711,605 | 12,334,147 | 1.89 | 93.6 |
| Doyogena | 35,184,652 | 12,256,756 | 2.07 | 93.8 |
| ShubiGemo | 33,804,134 | 12,376,058 | 1.98 | 93.6 |
| Loya | 33,822,572 | 12,172,853 | 1.98 | 93.6 |
| Hammari | 27,307,188 | 11,875,632 | 2.37 | 93.5 |
| Kabashi | 27,347,392 | 11,959,310 | 2.34 | 93.5 |
| Barbary | 36,190,249 | 12,695,368 | 2.31 | 93.7 |

**Table S4** InDel statistics for each sheep population

| **Population** | **Total number of InDels** | **No of Insertions** | **No of Deletions** | **dbSNP (%)** | **Average per sample** |
| --- | --- | --- | --- | --- | --- |
| Kefis | 4,102,070 | 1,782,561 | 2,319,509 | 93.9 | 1,388,390 |
| Segentu | 4,097,280 | 1,779,570 | 2,317,710 | 93.9 | 1,374,066 |
| Adane | 4,099,244 | 1,780,776 | 2,318,468 | 93.9 | 1,384,721 |
| Arabo | 4,100,430 | 1,781,492 | 2,318,938 | 94.1 | 1,390,390 |
| Gafera-Washera | 4,100,616 | 1,781,596 | 2,319,020 | 94 | 1,367,159 |
| Molale-Menz | 4,100,955 | 1,781,867 | 2,319,088 | 94 | 1,365,234 |
| Bonga | 4,099,254 | 1,780,829 | 2,318,425 | 93.9 | 1,341,735 |
| Gesses | 4,099,990 | 1,781,255 | 2,318,735 | 93.9 | 1,366,817 |
| Kido | 4,099,819 | 1,781,183 | 2,318,636 | 93.9 | 1,358,046 |
| Doyogena | 4,100,349 | 1,781,507 | 2,318,842 | 94 | 1,350,591 |
| ShubiGemo | 4,100,213 | 1,781,428 | 2,318,785 | 93.9 | 1,362,645 |
| Loya | 4,099,535 | 1,780,952 | 2,318,583 | 93.9 | 1,340,999 |
| Hammari | 3,496,053 | 1,657,173 | 1,838,880 | 93.7 | 1,622,816 |
| Kabashi | 3,650,551 | 1,730,407 | 1,920,144 | 93.6 | 1,598,529 |
| Barbary | 4,103,108 | 1,783,169 | 2,319,939 | 94 | 1,403,320 |

**Table S5** Number of SNPs in each category (150 sheep samples)

| **Category** | | **Number of SNPs** |
| --- | --- | --- |
| Total | | 34,857,882 |
| Upstream | | 1,093,873 |
| Exonic | Missense | 140,120 |
|  | Stop-gain | 3,374 |
|  | Stop-loss | 681 |
|  | Start-loss | 522 |
|  | Stop retained | 143 |
|  | Synonymous | 131,158 |
| Intronic | | 12,829,358 |
| Coding sequence | | 147 |
| 5_prime UTR | | 22,392 |
| 3_prime UTR | | 125,545 |
| Non coding transcript | | 24 |
| Non coding transcript exon | | 56,561 |
| Mature miRNA | | 247 |
| Splicing | | 25,838 |
| Splice donor | | 1,436 |
| Incomplete terminal codon | | 9 |
| Downstream | | 945,893 |
| Intergenic | | 19,479,560 |

**Table S6** Summary of average linkage disequilibrium (r2) values at different distances (Mb) across the 26 autosomes for the sheep groups.

| **CHR** | **ET-G1** | | **ET-G2** | | **SD** | | **LB** | |
| --- | --- | --- | --- | --- | --- | --- | --- | --- |
|  | ***r^2^*** | **Distance (Mb)** | ***r^2^*** | **Distance (Mb)** | ***r^2^*** | **Distance (Mb)** | ***r^2^*** | **Distance (Mb)** |
| 1 | 0.064 | 0.224 | 0.064 | 0.224 | 0.087 | 0.224 | 0.133 | 0.224 |
| 2 | 0.067 | 0.23 | 0.065 | 0.227 | 0.092 | 0.227 | 0.132 | 0.224 |
| 3 | 0.064 | 0.235 | 0.062 | 0.233 | 0.089 | 0.238 | 0.126 | 0.234 |
| 4 | 0.065 | 0.212 | 0.062 | 0.212 | 0.087 | 0.211 | 0.129 | 0.212 |
| 5 | 0.062 | 0.211 | 0.06 | 0.211 | 0.085 | 0.21 | 0.13 | 0.212 |
| 6 | 0.07 | 0.197 | 0.068 | 0.209 | 0.092 | 0.198 | 0.129 | 0.203 |
| 7 | 0.059 | 0.215 | 0.0578 | 0.206 | 0.082 | 0.214 | 0.125 | 0.212 |
| 8 | 0.065 | 0.204 | 0.06 | 0.219 | 0.087 | 0.202 | 0.123 | 0.207 |
| 9 | 0.069 | 0.195 | 0.064 | 0.208 | 0.09 | 0.191 | 0.126 | 0.198 |
| 10 | 0.083 | 0.197 | 0.083 | 0.195 | 0.101 | 0.191 | 0.143 | 0.189 |
| 11 | 0.062 | 0.213 | 0.064 | 0.215 | 0.086 | 0.218 | 0.133 | 0.218 |
| 12 | 0.067 | 0.195 | 0.062 | 0.201 | 0.087 | 0.202 | 0.127 | 0.205 |
| 13 | 0.066 | 0.215 | 0.06 | 0.209 | 0.085 | 0.215 | 0.129 | 0.217 |
| 14 | 0.059 | 0.223 | 0.053 | 0.206 | 0.085 | 0.21 | 0.125 | 0.211 |
| 15 | 0.068 | 0.192 | 0.065 | 0.199 | 0.09 | 0.198 | 0.143 | 0.202 |
| 16 | 0.065 | 0.189 | 0.063 | 0.188 | 0.085 | 0.187 | 0.124 | 0.191 |
| 17 | 0.064 | 0.194 | 0.064 | 0.188 | 0.087 | 0.203 | 0.125 | 0.200 |
| 18 | 0.062 | 0.201 | 0.062 | 0.199 | 0.083 | 0.197 | 0.137 | 0.199 |
| 19 | 0.062 | 0.206 | 0.058 | 0.196 | 0.089 | 0.201 | 0.128 | 0.214 |
| 20 | 0.064 | 0.190 | 0.062 | 0.190 | 0.087 | 0.201 | 0.132 | 0.190 |
| 21 | 0.069 | 0.177 | 0.065 | 0.192 | 0.087 | 0.182 | 0.129 | 0.185 |
| 22 | 0.069 | 0.186 | 0.064 | 0.193 | 0.089 | 0.175 | 0.134 | 0.181 |
| 23 | 0.065 | 0.188 | 0.065 | 0.189 | 0.085 | 0.185 | 0.129 | 0.183 |
| 24 | 0.056 | 0.187 | 0.056 | 0.197 | 0.084 | 0.194 | 0.123 | 0.201 |
| 25 | 0.063 | 0.176 | 0.059 | 0.184 | 0.083 | 0.173 | 0.123 | 0.178 |
| 26 | 0.064 | 0.179 | 0.061 | 0.18 | 0.084 | 0.19 | 0.132 | 0.186 |
| **Mean** | **0.065** | **0.201** | **0.063** | **0.203** | **0.087** | **0.201** | **0.13** | **0.203** |

**Table S7** The average autosomal effective population sizes over time for different sheep groups

| **Sheep Group** | **Generations ago** | | | | |
| --- | --- | --- | --- | --- | --- |
|  | **50** | **200** | **350** | **500** | **1000** |
| **ET-G1** | 1300 | 3550 | 3850 | 3750 | 2380 |
| **ET-G2** | 1500 | 3650 | 3850 | 3500 | 2680 |
| **SD** | 700 | 2300 | 3200 | 3242 | 2720 |
| **LB** | 350 | 700 | 1100 | 1395 | 1712 |

**Table S8** The candidate regions associated with high altitude adaptations detected by all methodologies by contrasting the ET-G2 Ethiopian sheep group vs. the LB and SD sheep

| **Reg.** | **Chr.** | **Start** | **Stop** | **Size (Mb)** | **No of**  **Genes** | **Genes** |
| --- | --- | --- | --- | --- | --- | --- |
| 1 | 1 | 19080001 | 19270000 | 0.190 | 11 | *RNF220, TMEM53, ARMH1, U5, KIF2C, RPS8, BEST4, PLK3, TCTEX1D4, BTBD19, U5* |
| 2 | 1 | 68600001 | 68840000 | 0.240 | 5 | *BTBD8, U6, C1orf146, GLMN, RPAP2* |
| 3 | 1 | 109920001 | 110020000 | 0.100 | 1 | *CD84* |
| 4 | 1 | 122430001 | 122620000 | 0.190 | 1 | *TIAM1* |
| 5 | 1 | 154650001 | 154790000 | 0.140 | 2 | *ZNF654, C3orf38* |
| 6 | 1 | 184180001 | 184370000 | 0.190 | 3 | *STXBP5L, POLQ, ARGFX* |
| 7 | 1 | 247990001 | 248180000 | 0.190 | 1 | *PRR23B* |
| 8 | 1 | 253550001 | 253740000 | 0.190 | 3 | *RAB6B, SRPRB, TF* |
| 9 | 2 | 1640001 | 1830000 | 0.190 | 2 | *EIPR1, TRAPPC12* |
| 10 | 2 | 11910001 | 12220000 | 0.310 | 2 | *ECPAS, OR2K2* |
| 11 | 2 | 68440001 | 68570000 | 0.130 | 1 | *KANK1* |
| 12 | 2 | 71850001 | 72040000 | 0.190 | 1 | *GLIS3* |
| 13 | 2 | 122520001 | 122620000 | 0.100 | 1 | *FSIP2* |
| 14 | 2 | 232440001 | 232900000 | 0.460 | 6 | *PTMA, PTPN4, PDE6D, COPS7B, SNORA62, DIS3L2* |
| 15 | 2 | 246240001 | 246430000 | 0.190 | 2 | *EMC1, UBR4* |
| 16 | 3 | 10630001 | 10880000 | 0.250 | 4 | *GAPVD1, HSPA5, RABEPK, PPP6C* |
| 17 | 3 | 10960001 | 11320000 | 0.360 | 10 | *SCAI, GOLGA1, ARPC5L, WDR38, U6, OLFML2A, oar-mir-181a-2, NR6A1, NR5A1, ADGRD2* |
| 18 | 3 | 11470001 | 11710000 | 0.240 | 1 | *LHX2* |
| 19 | 3 | 12160001 | 12370000 | 0.210 | 1 | *CRB2* |
| 20 | 3 | 77920001 | 78110000 | 0.190 | 2 | *EPAS1, PRKCE* |
| 21 | 3 | 90130001 | 90320000 | 0.190 | 2 | *FAM98A, RASGRP3* |
| 22 | 3 | 90320001 | 90580000 | 0.260 | 1 | *LTBP1* |
| 23 | 3 | 115160001 | 115350000 | 0.190 | 1 | *SYT1* |
| 24 | 3 | 154100001 | 154390000 | 0.290 | 1 | *MSRB3* |
| 25 | 3 | 174790001 | 174920000 | 0.130 | 1 | *POLR3B* |
| 26 | 3 | 180820001 | 180920000 | 0.100 | 2 | *USP18, ALG10* |
| 27 | 3 | 215580001 | 215680000 | 0.100 | 2 | *TNRC6B, ADSL* |
| 28 | 4 | 23770001 | 23870000 | 0.100 | 1 | *AGMO* |
| 29 | 4 | 73080001 | 73320000 | 0.240 | 1 | *ZNF804B* |
| 30 | 4 | 78890001 | 79010000 | 0.120 | 1 | *GLI3* |
| 31 | 4 | 96560001 | 96700000 | 0.140 | 1 | *CHCHD3* |
| 32 | 4 | 101290001 | 101400000 | 0.110 | 2 | *CREB3L2, 7SK* |
| 33 | 5 | 28800001 | 28930000 | 0.130 | 1 | *SNCAIP* |
| 34 | 5 | 41480001 | 41660000 | 0.180 | 5 | *ATP8B3, REXO1, ABHD17A, SCAMP4, CSNK1G2* |
| 35 | 5 | 48990001 | 49150000 | 0.160 | 5 | *5S_rRNA, U6, SRA1, APBB3, , SLC35A4* |
| 36 | 5 | 51690001 | 51880000 | 0.190 | 1 | *NR3C1* |
| 37 | 5 | 59560001 | 59700000 | 0.140 | 3 | *RBM22, DCTN4, SMIM3* |
| 38 | 5 | 65920001 | 66110000 | 0.190 | 2 | *HAVCR2, MED7* |
| 39 | 6 | 24710001 | 24890000 | 0.180 | 3 | *H2AZ1, DNAJB14, LAMTOR3* |
| 40 | 6 | 57900001 | 58090000 | 0.190 | 5 | *KLF3, TLR10, TLR1, TLR6, FAM114A1* |
| 41 | 6 | 69010001 | 69160000 | 0.150 | 1 | *LNX1* |
| 42 | 6 | 90570001 | 90760000 | 0.190 | 3 | *ART3, CXCL11, NUP54* |
| 43 | 6 | 116410001 | 116720000 | 0.310 | 7 | *CPLX1, DGKQ, FGFRL1, GAK, IDUA, SLC26A1, TMEM175* |
| 44 | 6 | 116830001 | 117040000 | 0.210 | 5 | *ATP5I, MFSD7, PCGF3, PDE6B, PIGG* |
| 45 | 7 | 14650001 | 14810000 | 0.160 | 2 | *PIAS1, CALML4* |
| 46 | 7 | 17340001 | 17530000 | 0.190 | 2 | *UACA, LARP6* |
| 47 | 8 | 12390001 | 12580000 | 0.190 | 3 | *TRMT11, HINT3, NCOA7* |
| 48 | 8 | 32040001 | 32230000 | 0.190 | 1 | *LIN28B* |
| 49 | 8 | 87360001 | 87470000 | 0.110 | 2 | *PDE10A, U6* |
| 50 | 9 | 28370001 | 28490000 | 0.120 | 5 | *MTSS1, NDUFB9, TATDN1, RNF139* |
| 51 | 9 | 28520001 | 28620000 | 0.100 | 1 | *TMEM65* |
| 52 | 9 | 36180001 | 36280000 | 0.100 | 3 | *PLAG1, CHCHD7, SDR16C5* |
| 53 | 9 | 67440001 | 67540000 | 0.100 | 2 | *SYBU, EBAG9* |
| 54 | 9 | 77030001 | 77200000 | 0.170 | 2 | *VPS13B, SNORA70* |
| 55 | 10 | 28560001 | 28820000 | 0.260 | 2 | *PDS5B, N4BP2L2* |
| 56 | 10 | 29390001 | 29510000 | 0.120 | 1 | *RXFP2* |
| 57 | 10 | 61190001 | 61380000 | 0.190 | 1 | *SLITRK6* |
| 58 | 11 | 18230001 | 18520000 | 0.290 | 4 | *NF1,EVI2A , EVI2B, OMG* |
| 59 | 11 | 37240001 | 37430000 | 0.190 | 6 | *CALCOCO2, TTLL6, HOXB13, HOXB9,HOXB8 , HOXB7* |
| 60 | 12 | 68390001 | 68510000 | 0.120 | 1 | *RPS6KC1* |
| 61 | 12 | 75850001 | 76040000 | 0.190 | 1 | *MIR181A-1* |
| 62 | 12 | 78290001 | 78440000 | 0.150 | 2 | *UBE2T, LGR6* |
| 63 | 13 | 38560001 | 38890000 | 0.330 | 4 | *RIN2, NAA20, CRNKL1, CFAP61* |
| 64 | 13 | 50470001 | 50650000 | 0.180 | 2 | *RNF24, PANK2* |
| 65 | 13 | 53340001 | 53660000 | 0.320 | 16 | *ZBTB46, ZGPAT, ARFRP1, TNFRSF6B, TNFRSF6B, STMN3, GMEB2, FNDC11, SRMS, PTK6, EEF1A2, KCNQ2, CHRNA4, ARFGAP1, BIRC7, YTHDF1* |
| 66 | 13 | 69250001 | 69350000 | 0.100 | 1 | *TOP1* |
| 67 | 13 | 69380001 | 69520000 | 0.140 | 2 | *PLCG1, ZHX3* |
| 68 | 15 | 21740001 | 21930000 | 0.190 | 6 | *DIXDC1,DLAT , PIH1D2, NKAPD1, TIMM8B, IL18* |
| 69 | 15 | 35110001 | 35280000 | 0.170 | 2 | *PLEKHA7, C11orf58* |
| 70 | 16 | 18150001 | 18290000 | 0.140 | 1 | *SMIM15* |
| 71 | 16 | 31620001 | 31810000 | 0.190 | 2 | *SEPP1, CCDC152* |
| 72 | 17 | 470001 | 660000 | 0.190 | 1 | *CPE* |
| 73 | 17 | 13200001 | 13340000 | 0.140 | 1 | *HHIP* |
| 74 | 17 | 34210001 | 34370000 | 0.160 | 1 | *FGF2* |
| 75 | 17 | 40470001 | 40660000 | 0.190 | 3 | *TMEM144, GASK1B, 5S_rRNA* |
| 76 | 18 | 5540001 | 5710000 | 0.170 | 1 | *ATP10A, ADAMTS17* |
| 77 | 18 | 32230001 | 32400000 | 0.170 | 5 | *PTPN9, SIN3A, MAN2C1, NEIL1, COMMD4* |
| 78 | 18 | 44550001 | 44740000 | 0.190 | 2 | *U6, RALGAPA1* |
| 79 | 19 | 550001 | 740000 | 0.190 | 2 | *VOPP1, LANCL2* |
| 80 | 19 | 23180001 | 23290000 | 0.110 | 1 | *CNTN4* |
| 81 | 19 | 30490001 | 30640000 | 0.150 | 1 | *FOXP1* |
| 82 | 19 | 43420001 | 43530000 | 0.110 | 1 | *SLMAP* |
| 83 | 19 | 51500001 | 51680000 | 0.180 | 7 | *NME6, CATHL1B , CATHL3, BAC5, SC5, CDC25A, 5S_rRNA* |
| 84 | 20 | 17230001 | 17480000 | 0.250 | 6 | *MAD2L1BP, 5S_rRNA, RSPH9, MRPS18A, , VEGFA* |
| 85 | 20 | 34130001 | 34320000 | 0.190 | 1 | *PRL* |
| 86 | 21 | 17690001 | 17880000 | 0.190 | 3 | *RSF1,CLNS1A , AQP11* |
| 87 | 22 | 10720001 | 10910000 | 0.190 | 6 | *LIPA, IFIT2, IFIT3, IFIT1, IFIT5, SLC16A12* |
| 88 | 22 | 50510001 | 50760000 | 0.250 | 4 | *MTG1, CYP2E1, ECHS1, FUOM* |
| 89 | 23 | 26220001 | 26380000 | 0.160 | 3 | *DSC1, DSC2, DSC3* |
| 90 | 24 | 9780001 | 9950000 | 0.170 | 3 | *CLEC16A, TNP2, PRM3* |
| 91 | 25 | 4050001 | 4180000 | 0.130 | 4 | *GNPAT, EXOC8, SPRTN, EGLN1* |
|  | | | | | **250** |  |

**Table S9** Enriched functional term clusters and their enrichment scores following DAVID and KOBAS analysis for genes identified in association with the study high-altitude environmental.

| ***Database*** | ***ID*** | ***Term*** | ***Gene Count*** | ***P-value*** | ***Genes*** | ***Benjamini*** |
| --- | --- | --- | --- | --- | --- | --- |
| KEGG | oas05152 | Tuberculosis | 8 | 0.000206 | *IL18, PLK3, TLR1, CATHL3, CALML4, BAC5, TLR6, SC5* | 0.039742 |
| GOTERM_BP | GO:0001666 | response to hypoxia | 6 | 0.000225 | *EPAS1, EGLN1, CHRNA4, VEGFA, NF1, ADSL* | 0.022153 |
| KEGG | oas04066 | HIF-1 signaling pathway | 4 | 0.001032 | *TF, EGLN1, VEGFA, PLCG1* | 0.124900 |
| KEGG | oas05200 | Pathways in cancer | 8 | 0.001061 | *EPAS1, EGLN1, RASGRP3, CALML4, PLCG1, VEGFA, FGF2, HHIP* | 0.102348 |
| GOTERM_BP | GO:0045766 | positive regulation of angiogenesis | 2 | 0.001288 | *VEGFA, FGF2* | 0.137436 |
| KEGG | oas04014 | Ras signaling pathway | 7 | 0.002274 | *RASGRP3, CALML4, TIAM1, PLCG1, VEGFA, NF1, FGF2* | 0.146315 |
| KEGG | oas01521 | EGFR tyrosine kinase inhibitor resistance | 4 | 0.003660 | *PLCG1, VEGFA, FGF2, NF1* | 0.162392 |
| KEGG | oas00230 | Purine metabolism | 5 | 0.004207 | *ADSL, PDE10A, POLR3B, NME6, PDE6D, PDE6B* | 0.162392 |
| KEGG | oas04015 | Rap1 signaling pathway | 6 | 0.005478 | *RASGRP3, CALML4, TIAM1, PLCG1, VEGFA, FGF2* | 0.176193 |
| KEGG | oas05211 | Renal cell carcinoma | 3 | 0.005766 | *EPAS1, EGLN1, VEGFA* | 0.109797 |
| KEGG | oas04970 | Salivary secretion | 4 | 0.006879 | *SC5, BAC5, CATHL3, CALML4* | 0.189659 |
| GOTERM_BP | GO:0050707 | regulation of cytokine secretion | 3 | 0.008010 | *TLR1, TLR10, TLR6* | 1.000000 |
| KEGG | oas03060 | Protein export | 2 | 0.012862 | *SRPRB, HSPA5* | 0.275811 |

**Table S10** The candidate regions associated with arid/desert low-altitude environment detected via ZHp of LB and SD sheep using all methodologies by contrasting the LB and SD sheep groups vs. the ET-G2 Ethiopian sheep from high-altitude African environments.

| **Reg.** | **Chr.** | **Start** | **Stop** | **Size (Mb)** | **No of**  **Genes** | **Genes** |
| --- | --- | --- | --- | --- | --- | --- |
| 1 | 1 | 68600001 | 68840000 | 0.240 | 5 | *BTBD8, U6, C1orf146, GLMN, RPAP2* |
| 2 | 1 | 90180001 | 90370000 | 0.190 | 6 | *PHTF1, RSBN1, PTPN22, BCL2L15, AP4B1, DCLRE1B* |
| 3 | 1 | 109890001 | 110040000 | 0.150 | 1 | *CD84, SLAMF1* |
| 4 | 1 | 119540001 | 119670000 | 0.130 | 2 | *MRPS6, SLC5A3* |
| 5 | 1 | 129020001 | 129140000 | 0.120 | 2 | *MRPL39, U6* |
| 6 | 1 | 154650001 | 154790000 | 0.140 | 2 | *ZNF654, C3orf38* |
| 7 | 1 | 237090001 | 237200000 | 0.110 | 3 | *HPS3, HLTF, GYG1* |
| 8 | 1 | 253550001 | 253740000 | 0.190 | 3 | *RAB6B, SRPRB, TF* |
| 9 | 2 | 8860001 | 9050000 | 0.190 | 1 | *TNFSF15* |
| 10 | 2 | 43120001 | 43310000 | 0.190 | 10 | *PHYHIP, BMP1, SFTPC, LGI3, REEP4, HR, NUDT18, FAM160B2, DMTN, FGF17* |
| 11 | 2 | 52290001 | 52540000 | 0.250 | 11 | *TMEM8B, FAM221B, HINT2, SPAG8, NPR2, MSMP, RGP1, GBA2, CREB3, TLN1, TPM2* |
| 12 | 2 | 58610001 | 58760000 | 0.150 | 2 | *GNAQ* |
| 13 | 2 | 68440001 | 68570000 | 0.130 | 1 | *KANK1* |
| 14 | 2 | 117950001 | 118090000 | 0.140 | 2 | *HIBCH, C2orf88* |
| 15 | 2 | 120860001 | 121050000 | 0.190 | 1 | *CALCRL* |
| 16 | 2 | 121340001 | 121440000 | 0.100 | 1 | *ZSWIM2* |
| 17 | 2 | 122520001 | 122620000 | 0.100 | 1 | *FSIP2* |
| 18 | 2 | 174920001 | 175040000 | 0.120 | 1 | *TMEM163* |
| 19 | 2 | 184320001 | 184480000 | 0.160 | 4 | *EPB41L5, U4, TMEM185B, RALB* |
| 20 | 2 | 210250001 | 210440000 | 0.190 | 2 | *MAP2, UNC80* |
| 21 | 2 | 219370001 | 219510000 | 0.140 | 5 | *SLC11A1, CTDSP1, oar-mir-26b, VIL1, USP37* |
| 22 | 2 | 232440001 | 232630000 | 0.190 | 2 | *PDE6D, COPS7B* |
| 23 | 2 | 232660001 | 233000000 | 0.340 | 2 | *SNORA62, DIS3L2* |
| 24 | 2 | 234290001 | 234530000 | 0.240 | 13 | *BSDC1, TSSK3, FAM229A, HDAC1, MARCKSL1, LCK, FAM167B, DCDC2B, IQCC, TXLNA, KPNA6, U6, TMEM39B* |
| 25 | 3 | 10960001 | 11320000 | 0.360 | 9 | *GOLGA1, ARPC5L, , WDR38, U6, OLFML2A, oar-mir-181a-2, NR6A1, NR5A1, ADGRD2* |
| 26 | 3 | 11470001 | 11710000 | 0.240 | 1 | *LHX2* |
| 27 | 3 | 12160001 | 12410000 | 0.250 | 1 | *CRB2* |
| 28 | 3 | 45050001 | 45160000 | 0.110 | 2 | *OTX1, EHBP1* |
| 29 | 3 | 129680001 | 129840000 | 0.160 | 2 | *SOCS2, CRADD* |
| 30 | 3 | 154100001 | 154390000 | 0.290 | 1 | *MSRB3* |
| 31 | 3 | 174810001 | 174920000 | 0.110 | 1 | *POLR3B* |
| 32 | 3 | 180820001 | 180920000 | 0.100 | 2 | *USP18, ALG10* |
| 33 | 3 | 202350001 | 202540000 | 0.190 | 2 | *LRP6, MANSC1* |
| 34 | 4 | 23770001 | 23870000 | 0.100 | 1 | *AGMO* |
| 35 | 4 | 48530001 | 48750000 | 0.220 | 6 | *COG5, GPR22, DUS4L, BCAP29, SLC26A4* |
| 36 | 4 | 68650001 | 68840000 | 0.190 | 1 | *EVX1* |
| 37 | 4 | 87490001 | 87680000 | 0.190 | 1 | *SLC13A1* |
| 38 | 4 | 96560001 | 96700000 | 0.140 | 1 | *CHCHD3* |
| 39 | 4 | 101440001 | 101620000 | 0.180 | 3 | *CREB3L2, AKR1D1, TRIM24* |
| 40 | 5 | 6330001 | 6430000 | 0.100 | 1 | *EPS15L1* |
| 41 | 5 | 15510001 | 15700000 | 0.190 | 13 | *ALKBH7, CD70, CLPP, CRB3, DENND1C, GTF2F1, KHSRP, PSPN, SLC25A23, SLC25A41, TNFSF9, TRNAV-CAC, TUBB4A* |
| 42 | 5 | 49090001 | 49190000 | 0.100 | 8 | *SRA1, APBB3, , SLC35A4, CD14, TMCO6, NDUFA2, IK, DND1, HARS1* |
| 43 | 5 | 49290001 | 49480000 | 0.190 | 3 | *PCDHA5, PCDHA11, PCDHAC2* |
| 44 | 5 | 51690001 | 51880000 | 0.190 | 1 | *NR3C1* |
| 45 | 5 | 58020001 | 58130000 | 0.110 | 1 | *ABLIM3* |
| 46 | 5 | 59560001 | 59700000 | 0.140 | 3 | *RBM22, DCTN4, SMIM3* |
| 47 | 6 | 19850001 | 20040000 | 0.190 | 1 | *PPA2* |
| 48 | 6 | 24660001 | 24880000 | 0.220 | 2 | *H2AZ1, DNAJB14* |
| 49 | 6 | 44720001 | 44940000 | 0.220 | 3 | *ZCCHC4, ANAPC4, 5S_rRNA* |
| 50 | 6 | 69060001 | 69160000 | 0.100 | 1 | *LNX1* |
| 51 | 6 | 69830001 | 70090000 | 0.260 | 1 | *FGF5* |
| 52 | 6 | 94560001 | 94750000 | 0.190 | 1 | *FGF5* |
| 53 | 6 | 103780001 | 103970000 | 0.190 | 1 | *MSX1* |
| 54 | 6 | 116410001 | 117040000 | 0.630 | 7 | *GAK, CPLX1, UVSSA, MAEA, SLC49A3, PDE6B, PIGG* |
| 55 | 7 | 51260001 | 51450000 | 0.190 | 2 | *RFX7, NEDD4* |
| 56 | 7 | 55870001 | 56140000 | 0.270 | 1 | *CYP19* |
| 57 | 7 | 57170001 | 57310000 | 0.140 | 1 | *ATP8B4* |
| 58 | 8 | 32040001 | 32230000 | 0.190 | 1 | *LIN28B* |
| 59 | 8 | 62710001 | 62810000 | 0.100 | 1 | *TNFAIP3, 7SK* |
| 60 | 9 | 28440001 | 28540000 | 0.100 | 2 | *TATDN1, RNF139* |
| 61 | 9 | 36180001 | 36280000 | 0.100 | 3 | *PLAG1, CHCHD7, SDR16C5* |
| 62 | 10 | 28560001 | 28820000 | 0.260 | 2 | *PDS5B, N4BP2L2* |
| 63 | 10 | 29390001 | 29510000 | 0.120 | 2 | *RXFP2* |
| 64 | 10 | 36140001 | 36420000 | 0.280 | 5 | *GJB6, GJB2, GJA3, ZMYM2, CRYL1* |
| 65 | 11 | 10750001 | 10940000 | 0.190 | 2 | *BRIP1, INTS2* |
| 66 | 11 | 18170001 | 18520000 | 0.350 | 4 | *NF1,EVI2A , EVI2B, OMG* |
| 67 | 11 | 18820001 | 19010000 | 0.190 | 2 | *KSR1, NOS2* |
| 68 | 11 | 26350001 | 26540000 | 0.190 | 8 | *ALOX12, ASGR1, ASGR2, BCL6B, C11H17orf49, RNASEK, SLC16A11, SLC16A13* |
| 69 | 11 | 27010001 | 27200000 | 0.190 | 8 | *CHD3, CNTROB, DNAH2, KCNAB3, KDM6B, NAA38, TMEM88, TRAPPC1* |
| 70 | 11 | 51270001 | 51380000 | 0.110 | 1 | *RNF213* |
| 71 | 12 | 27770001 | 27870000 | 0.100 | 1 | *ITPKB* |
| 72 | 12 | 42800001 | 42900000 | 0.100 | 1 | *RERE* |
| 73 | 12 | 68390001 | 68510000 | 0.120 | 1` | *RPS6KC1* |
| 74 | 12 | 75860001 | 75990000 | 0.130 | 1 | *oar-mir-181a-1* |
| 75 | 12 | 78290001 | 78440000 | 0.150 | 2 | *U6, UBE2T,* |
| 76 | 12 | 78440001 | 78550000 | 0.110 | 1 | *CSRP1* |
| 77 | 12 | 78990001 | 79100000 | 0.110 | 3 | *RNPEP, ELF3, PTPN7* |
| 78 | 13 | 17150001 | 17250000 | 0.100 | 1 | *CREM* |
| 79 | 13 | 38560001 | 38890000 | 0.330 | 4 | *RIN2, NAA20, CRNKL1, CFAP61* |
| 80 | 13 | 46130001 | 46320000 | 0.190 | 4 | *PRNP, PRND, PRNT, RASSF2* |
| 81 | 13 | 50420001 | 50670000 | 0.250 | 2 | *RNF24, PANK2* |
| 82 | 13 | 53090001 | 53270000 | 0.180 | 10 | *NPBWR2, OPRL1, RGS19, TCEA2, PRPF6, U6, ZNF512B, UCKL1, DNAJC5, TPD52L2* |
| 83 | 13 | 53290001 | 53640000 | 0.350 | 6 | *ZBTB46, ZGPAT, ARFRP1, STMN3, GMEB2, FNDC11* |
| 84 | 13 | 69250001 | 69350000 | 0.100 | 1 | *TOP1* |
| 85 | 13 | 76710001 | 76900000 | 0.190 | 4 | *ARFGEF2, CSE1L, STAU1, U6* |
| 86 | 14 | 34380001 | 34590000 | 0.210 | 11 | *HSD11B2, ATP6V0D1, AGRP, RIPOR1, CTCF, CARMIL2, ACD, PARD6A, ENKD1, C16orf86, GFOD2* |
| 87 | 14 | 37450001 | 37570000 | 0.120 | 1 | *ZFHX3* |
| 88 | 14 | 48230001 | 48420000 | 0.190 | 8 | *SAMD4B, PAF1, MED29, ZFP36, PLEKHG2, SUPT5H, TIMM50, DLL3* |
| 89 | 14 | 49510001 | 49700000 | 0.190 | 9 | *ERICH4, CYP2B6, DMAC2, B3GNT8, BCKDHA, EXOSC5, , B9D2, TGFB1, CCDC97* |
| 90 | 15 | 28570001 | 28760000 | 0.190 | 7 | *UBE4A, SNORA70, TTC36, TMEM25, IFT46, ARCN1, PHLDB1* |
| 91 | 15 | 35110001 | 35280000 | 0.170 | 2 | *PLEKHA7, C11orf58* |
| 92 | 15 | 50900001 | 51090000 | 0.190 | 4 | *P2RY6, ARHGEF17, RELT, FAM168A* |
| 93 | 16 | 31650001 | 31750000 | 0.100 | 2 | *SEPP1, CCDC152* |
| 94 | 17 | 13130001 | 13340000 | 0.210 | 1 | *HHIP* |
| 95 | 17 | 34210001 | 34370000 | 0.160 | 1 | *FGF2* |
| 96 | 17 | 52360001 | 52570000 | 0.210 | 4 | *HCAR1, KNTC1, RSRC2, ZCCHC8,* |
| 97 | 17 | 60890001 | 61080000 | 0.190 | 3 | *TPCN1, SLC8B1, RPH3A* |
| 98 | 17 | 62000001 | 62270000 | 0.270 | 6 | *BICDL1, RAB35, GCN1, PXN, U4, SIRT4* |
| 99 | 18 | 5540001 | 5710000 | 0.170 | 1 | *ADAMTS17* |
| 100 | 18 | 20500001 | 20690000 | 0.190 | 2 | *ZNF710, IDH2* |
| 101 | 18 | 22650001 | 22790000 | 0.140 | 1 | *BNC1* |
| 102 | 18 | 23610001 | 23850000 | 0.240 | 1 | *MEX3B* |
| 103 | 18 | 32210001 | 32410000 | 0.200 | 5 | *PTPN9, SIN3A, MAN2C1, NEIL1, COMMD4* |
| 104 | 18 | 54230001 | 54420000 | 0.190 | 12 | *CATSPER2, CKMT1B, ELL3, HYPK, MAP1A, MFAP1, PDIA3, PPIP5K1, SERF2, SERINC4, STRC, WDR76* |
| 105 | 19 | 23200001 | 23420000 | 0.220 | 1 | *CNTN4* |
| 106 | 19 | 30490001 | 30640000 | 0.150 | 1 | *FOXP1* |
| 107 | 19 | 43420001 | 43530000 | 0.110 | 1 | *SLMAP* |
| 108 | 19 | 51500001 | 51680000 | 0.180 | 6 | *NME6, CATHL3, BAC5, SC5, CDC25A, 5S_rRNA* |
| 109 | 19 | 55290001 | 55480000 | 0.190 | 2 | *ATG7, VGLL4* |
| 110 | 20 | 15030001 | 15220000 | 0.190 | 8 | *UNC5CL, TSPO2, APOBEC2, OARD1, NFYA, , TREML1, TREM2, TREML2* |
| 111 | 20 | 17230001 | 17480000 | 0.250 | 5 | *MAD2L1BP, 5S_rRNA, RSPH9, MRPS18A, VEGFA* |
| 112 | 20 | 31990001 | 32180000 | 0.190 | 4 | *TDP2, KIAA0319, ALDH5A1, GPLD1* |
| 113 | 21 | 49760001 | 49890000 | 0.130 | 6 | *IFITM5, PGGHG, PSMD13, SIRT3, RIC8A, ODF3* |
| 114 | 22 | 8940001 | 9040000 | 0.100 | 1 | *MINPP1* |
| 115 | 22 | 50600001 | 50760000 | 0.160 | 2 | *CYP2E1, ECHS1* |
| 116 | 23 | 26220001 | 26380000 | 0.160 | 2 | *DSC1, DSC2* |
| 117 | 24 | 9780001 | 9950000 | 0.170 | 3 | *CLEC16A, TNP2, PRM3* |
| 118 | 24 | 14110001 | 14230000 | 0.120 | 2 | *MYH11, CEP20* |
| 119 | 24 | 35930001 | 36030000 | 0.100 | 7 | *NYAP1, TSC22D4, C7orf61, PPP1R35, MEPCE, ZCWPW1, U6* |
| 120 | 24 | 38210001 | 38400000 | 0.190 | 5 | *KDELR2, GRID2IP, ZDHHC4, C7orf26, ZNF12* |
| 121 | 24 | 41720001 | 41850000 | 0.130 | 1 | *FAM20C* |
| 122 | 25 | 4060001 | 4180000 | 0.120 | 4 | *GNPAT, EXOC8, SPRTN, EGLN1* |
| 123 | 25 | 17460001 | 17580000 | 0.120 | 1 | *C25H10orf107* |
|  |  |  |  |  | **182** |  |

**Table S11** Enriched functional term clusters and their enrichment scores following DAVID and KOBAS analysis for genes identified in association with the study low-altitude arid environmental challenges.

| ***Database*** | ***ID*** | ***Term*** | ***Gene Count*** | ***P-value*** | ***Genes*** | ***Benjamini*** |
| --- | --- | --- | --- | --- | --- | --- |
| GOTERM_MF | GO:0008270 | zinc ion binding | 36 | 0.0002 | *ZCCHC8, HHIP, PXN, TNFAIP3, CHD3, NR3C1, ZDHHC4, ZCWPW1, ZMYM2, RNF213, CSRP1, ABLIM3, RNF139, HLTF, RNPEP, ADAMTS17, TRIM24, MAN2C1, ZSWIM2, ZFHX3, APOBEC2, RNF24, SIRT4, SIRT3, NR5A1, LIN28B, MEX3B, NR6A1, BMP1, LHX2, LOC101120178, LNX1, TCEA2, MSRB3, NEIL1, ZCCHC4* | 0.050324 |
| GOTERM_CC | GO:0005634 | nucleus | 4 | 0.0023 | *ZGPAT, NR3C1, ZFP36, TGFB1* | 0.942708 |
| KEGG | oas04926 | Relaxin signaling pathway | 6 | 0.0024 | *TGFB1, NOS2, RXFP2, CREB3, CREB3L2, VEGFA* | 1.000000 |
| KEGG | oas04918 | Thyroid hormone synthesis | 6 | 0.0025 | *CREB3, GNAQ, CREB3L2, LOC101116157, SLC26A4, ASGR2* | 0.838712 |
| KEGG | oas04015 | Rap1 signaling pathway | 8 | 0.0030 | *RALB, GNAQ, PARD6A, TLN1, VEGFA, FGF5, FGF17, FGF2* | 1.000000 |
| KEGG | oas04925 | Aldosterone synthesis and secretion | 3 | 0.0031 | *CREB3L2, CREB3, GNAQ* | 1.000000 |
| KEGG | oas04962 | Vasopressin-regulated water reabsorption | 3 | 0.0036 | *CREB3L2, CREB3, DCTN4* | 1.000000 |
| GOTERM_CC | GO:0005922 | connexon complex | 3 | 0.0039 | *GJB2, GJA3, GJB6* | 1.000000 |
| GOTERM_CC | GO:0005737 | cytoplasm | 59 | 0.0045 | *GMEB2, CSE1L, BICDL1, TNFAIP3, SRA1, STMN3, PTPN22, CHD3, NR3C1, PHTF1, ZFP36, RASSF2, SIN3A, TRIM24, KPNA6, KNTC1, UBE4A, KCNAB3, PRM3, COMMD4, CDC25A, TXLNA, TLN1, NEIL1, IFT46, RBM22, NAA20, PXN, CREM, COPS7B, RPAP2, EHBP1, SOCS2, EPB41L5, BRIP1, MAP2, DND1, MSX1, ATG7, APBB3, GCN1, EGLN1, ACD, ZFHX3, TGFB1, SRMS, SPAG8, CRADD, SAMD4B, MAD2L1BP, PTK6, MSMP, PLEKHA7, TDP2, LNX1, NF1, PAF1, PTPN9* | 0.902817 |
| GOTERM_CC | GO:0005783 | endoplasmic reticulum | 14 | 0.0052 | *PDIA3, PRNP, CALCRL, STAU1, ARCN1, CREB3, HSD11B2, REEP4, AGMO, CREB3L2, BCAP29, MSRB3, CYP2E1, PIGG* | 1.000000 |
| KEGG | oas04927 | Cortisol synthesis and secretion | 4 | 0.0053 | *CREB3L2, CREB3, GNAQ, NR5A1* | 1.000000 |
| KEGG | oas05016 | Huntington disease | 8 | 0.0074 | *DNAH2, HDAC1, NDUFA2, GNAQ, CREB3, CREB3L2, DCTN4, SIN3A* | 1.000000 |
| KEGG | oas00640 | Propanoate metabolism | 3 | 0.0133 | *ECHS1, HIBCH, BCKDHA* | 1.000000 |
| KEGG | oas00140 | Steroid hormone biosynthesis | 4 | 0.0140 | *AKR1D1, HSD11B2, CYP2E1, CYP19* | 1.000000 |

**Table S12** The candidate regions associated with tail length as detected by all of three methods (*ZHp, XP-EHH, F_ST_*) of detecting selection signatures amongst two broad groups of sheep viz long tailed sheep (ET-G2 and SD) and short tailed sheep (ET-G1 and LB) analysed in this study.

| **Reg** | **Chr** | **Start** | **Stop** | **Size (Mb)** | **No**  **of**  **Genes** | **Genes** |
| --- | --- | --- | --- | --- | --- | --- |
| 1 | 1 | 5930001 | 6120000 | 0.190 | 0 | *-* |
| 2 | 1 | 68600001 | 68840000 | 0.240 | 4 | *BTBD8, C1orf146, GLMN, RPAP2* |
| 3 | 1 | 94840001 | 94980000 | 0.140 | 0 | *-* |
| 4 | 1 | 109860001 | 110040000 | 0.180 | 3 | *ENSOARG00000008861, CD84, SLAMF1* |
| 5 | 1 | 169610001 | 169800000 | 0.190 | 0 | *-* |
| 6 | 1 | 184180001 | 184370000 | 0.190 | 3 | *STXBP5L, POLQ, ENSOARG00000020026* |
| 7 | 1 | 253550001 | 253730000 | 0.180 | 4 | *RAB6B, SRPRB, TF, ENSOARG00000008517* |
| 8 | 2 | 1640001 | 1830000 | 0.190 | 3 | *ENSOARG00000025634, EIPR1, TRAPPC12* |
| 9 | 2 | 11910001 | 12100000 | 0.190 | 3 | *ENSOARG00000006672, ECPAS, OR2K2* |
| 10 | 2 | 52290001 | 52530000 | 0.240 | 10 | *TMEM8B, FAM221B, HINT2, SPAG8, NPR2, MSMP, RGP1, GBA2, CREB3, TLN1* |
| 11 | 2 | 68440001 | 68570000 | 0.130 | 1 | *KANK1* |
| 12 | 2 | 73560001 | 73750000 | 0.190 | 3 | *KIAA2026, ENSOARG00000013612, ENSOARG00000013618* |
| 13 | 2 | 122520001 | 122620000 | 0.100 | 2 | *FSIP2, ENSOARG00000025803* |
| 14 | 2 | 232440001 | 232630000 | 0.190 | 3 | *PDE6D, COPS7B, ENSOARG00000020723* |
| 15 | 2 | 232660001 | 232900000 | 0.240 | 2 | *SNORA62, DIS3L2* |
| 16 | 2 | 240710001 | 240890000 | 0.180 | 1 | *RUNX3* |
| 17 | 3 | 10630001 | 10820000 | 0.190 | 6 | *GAPVD1, HSPA5, RABEPK, PPP6C, SCAI, ENSOARG00000025028* |
| 18 | 3 | 10920001 | 11320000 | 0.400 | 11 | *GOLGA1, ARPC5L, ENSOARG00000013275, WDR38, OLFML2A, oar-mir-26a, ENSOARG00000023155, U6, NR5A1, NR6A1, ADGRD2* |
| 19 | 3 | 11190001 | 11300000 | 0.110 | 2 | *NR5A1, NR6A1* |
| 20 | 3 | 11760001 | 11950000 | 0.190 | 2 | *ENSOARG00000013754, ENSOARG00000019124* |
| 21 | 3 | 12220001 | 12410000 | 0.190 | 3 | *ENSOARG00000013769, ENSOARG00000021273, CRB2* |
| 22 | 3 | 90320001 | 90580000 | 0.260 | 3 | *RASGRP3, ENSOARG00000025063, LTBP1* |
| 23 | 3 | 154100001 | 154390000 | 0.290 | 1 | *MSRB3* |
| 24 | 3 | 161530001 | 161710000 | 0.180 | 12 | *CTDSP2, oar-mir-26a, AVIL, TSFM, EEF1AKMT3, METTL1, CYP27B1, MARCHF9, CDK4, TSPAN31, AGAP2, OS9* |
| 25 | 4 | 68010001 | 68180000 | 0.170 | 1 | *JAZF1* |
| 26 | 4 | 73080001 | 73270000 | 0.190 | 1 | *U6* |
| 27 | 4 | 96540001 | 96640000 | 0.100 | 1 | *CHCHD3* |
| 28 | 5 | 15510001 | 15700000 | 0.190 | 9 | *TUBB4A, DENND1C, CRB3, SLC25A23, SLC25A41, KHSRP, GTF2F1, ALKBH7, CLPP* |
| 29 | 5 | 51670001 | 51890000 | 0.220 | 2 | *ARHGAP26, NR3C1* |
| 30 | 5 | 107340001 | 107440000 | 0.100 | 1 | *WDR36* |
| 31 | 6 | 10070001 | 10260000 | 0.190 | 1 | *LOC105613665* |
| 32 | 6 | 57900001 | 58090000 | 0.190 | 5 | *TLR10, TLR1, TLR6, KLF3, FAM114A1* |
| 33 | 6 | 69010001 | 70090000 | 1.079 | 4 | *PDGFRA, LNX1, GSX2, CHIC2* |
| 34 | 6 | 115960001 | 117040000 | 1.079 | 5 | *LETM1, NELFA, NSD2, SLC49A3, PDE6B* |
| 35 | 7 | 55870001 | 56060000 | 0.190 | 3 | *NF1, EVI2A, EVI2B, OMG* |
| 36 | 7 | 56080001 | 56300000 | 0.220 | 2 | *CYP19, ENSOARG00000000908* |
| 37 | 8 | 11480001 | 11620000 | 0.140 | 0 | *-* |
| 38 | 8 | 32040001 | 32230000 | 0.190 | 1 | *LIN28B* |
| 39 | 8 | 51540001 | 51670000 | 0.130 | 0 | *-* |
| 40 | 10 | 7310001 | 7500000 | 0.190 | 1 | *ENSOARG00000006641* |
| 41 | 10 | 28560001 | 28820000 | 0.260 | 2 | *PDS5B, N4BP2L2* |
| 42 | 10 | 29350001 | 29540000 | 0.190 | 4 | *ENSOARG00000011616, ENSOARG00000011616, RXFP2* |
| 43 | 10 | 80430001 | 80620000 | 0.190 | 0 | *-* |
| 44 | 11 | 10750001 | 10940000 | 0.190 | 2 | *INTS2, BRIP1* |
| 45 | 11 | 18230001 | 18520000 | 0.290 | 4 | *NF1, EVI2A, EVI2B, OMG* |
| 46 | 11 | 26350001 | 26540000 | 0.190 | 9 | *ALOX12, RNASEK, U6, BCL6B, SLC16A13, SLC16A11, CLEC10A, ASGR2, ASGR1* |
| 47 | 11 | 27010001 | 27200000 | 0.190 | 10 | *DNAH2, NAA38, KDM6B, TMEM88, CYB5D1, CHD3, RNF227, KCNAB3, TRAPPC1, CNTROB* |
| 48 | 11 | 37200001 | 37430000 | 0.230 | 7 | *CALCOCO2, TTLL6, HOXB13, HOXB9, ENSOARG00000026370, HOXB8, HOXB7* |
| 49 | 12 | 75850001 | 76040000 | 0.190 | 2 | *ENSOARG00000021991, oar-mir-181a-1* |
| 50 | 13 | 38560001 | 38880000 | 0.320 | 4 | *RIN2, NAA20, CRNKL1, CFAP61* |
| 51 | 13 | 42670001 | 42800000 | 0.130 | 3 | *LOC10110963 (dihydrodiol dehydrogenase 3 like), LOC101109377 (dihydrodiol dehydrogenase 3 like), LOC101109111 (dihydrodiol dehydrogenase 3)* |
| 52 | 13 | 53450001 | 53640000 | 0.190 | 8 | *ENSOARG00000026240, SRMS, PTK6, ENSOARG00000010995, EEF1A2, KCNQ2, CHRNA4, ARFGAP1,* |
| 53 | 13 | 62990001 | 63130000 | 0.140 | 3 | *SNORA73, ENSOARG00000009308, ENSOARG00000009354* |
| 54 | 14 | 48390001 | 48580000 | 0.190 | 4 | *ENSOARG00000006291, DYRK1B, FBL, ENSOARG00000006366* |
| 55 | 15 | 21740001 | 21930000 | 0.190 | 7 | *DIXDC1, ENSOARG00000015712, PIH1D2, NKAPD1, TIMM8B, ENSOARG00000016150, IL18* |
| 56 | 15 | 56010001 | 56220000 | 0.210 | 4 | *U6, ENSOARG00000026835, ENSOARG00000006821, CCDC34* |
| 57 | 15 | 72470001 | 72620000 | 0.150 | 3 | *EXT2, U6, ALX4* |
| 58 | 15 | 77750001 | 77950000 | 0.200 | 11 | *ENSOARG00000008882, SSRP1, P2RX3, SLC43A1* |
| 59 | 16 | 31620001 | 31810000 | 0.190 | 2 | *ENSOARG00000008775, CCDC152* |
| 60 | 17 | 13210001 | 13330000 | 0.120 | 1 | *HHIP* |
| 61 | 17 | 29820001 | 30220000 | 0.400 | 1 | *ENSOARG00000021611* |
| 62 | 18 | 50910001 | 51100000 | 0.190 | 0 | *-* |
| 63 | 19 | 23200001 | 23420000 | 0.220 | 1 | *CNTN4* |
| 64 | 20 | 17230001 | 17480000 | 0.250 | 6 | *MAD2L1BP, 5S_rRNA, RSPH9, MRPS18A, ENSOARG00000008711, VEGFA* |
| 65 | 20 | 25450001 | 25590000 | 0.140 | 3 | *ENSOARG00000015660, ENSOARG00000015707, ENSOARG00000015866* |
| 66 | 23 | 1030001 | 1220000 | 0.190 | 1 | *ENSOARG00000026145* |
| 67 | 25 | 4050001 | 4160000 | 0.110 | 4 | *GNPAT, EXOC8, SPRTN, EGLN1* |
| 68 | 1 | 5930001 | 6120000 | 0.190 | 0 | *-* |
| **Total** | | | | | **225** |  |

**Table S13** Candidate regions associated with tail fat-depots as detected by all methodologies (ZHp, XP-EHH, F_ST_) of detecting selection signatures as employed in the current study.

| **Reg.** | **Chr.** | **Start** | **Stop** | **Size (Mb)** | **No of Genes** | **Genes** |
| --- | --- | --- | --- | --- | --- | --- |
| 1 | 1 | 68690001 | 68860000 | 0.170 | 3 | *C1orf146, GLMN, RPAP2* |
| 2 | 2 | 52290001 | 52540000 | 0.250 | 11 | *TMEM8B, FAM221B, HINT2, SPAG8, NPR2, MSMP, RGP1, GBA2, CREB3, TLN1, TPM2* |
| 3 | 2 | 73490001 | 73640000 | 0.150 | 3 | *MLANA, KIAA2026, ENSOARG00000013612* |
| 4 | 2 | 122330001 | 122530000 | 0.200 | 2 | *ENSOARG00000016633, FSIP2* |
| 5 | 2 | 210250001 | 210440000 | 0.190 | 2 | *ENSOARG00000019133, MAP2* |
| 6 | 2 | 232660001 | 232870000 | 0.210 | 2 | *SNORA62, DIS3L2* |
| 7 | 2 | 240710001 | 240890000 | 0.180 | 1 | *RUNX3* |
| 8 | 3 | 10650001 | 10810000 | 0.160 | 5 | *GAPVD1, HSPA5, RABEPK, PPP6C, SCAI* |
| 9 | 3 | 11760001 | 11950000 | 0.190 | 2 | *ENSOARG00000013754, ENSOARG00000019124* |
| 10 | 3 | 12160001 | 12370000 | 0.210 | 3 | *CRB2, ENSOARG00000013754, ENSOARG00000013769* |
| 11 | 3 | 124800001 | 124920000 | 0.120 | 0 | *-* |
| 12 | 3 | 154130001 | 154350000 | 0.220 | 1 | *MSRB3* |
| 13 | 4 | 69930001 | 70070000 | 0.140 | 1 | *ENSOARG00000025241* |
| 14 | 4 | 87490001 | 87680000 | 0.190 | 1 | *SLC13A1* |
| 15 | 5 | 15510001 | 15700000 | 0.190 | 9 | *TUBB4A, DENND1C, CRB3, SLC25A23, SLC25A41, KHSRP, GTF2F1, ALKBH7, CLPP* |
| 16 | 5 | 107140001 | 107360000 | 0.220 | 1 | *ENSOARG00000010481* |
| 17 | 6 | 10070001 | 10260000 | 0.190 | 1 | *LOC105613665* |
| 18 | 6 | 36140001 | 36430000 | 0.290 | 5 | *ENSOARG00000000388, ENSOARG00000000447, HERC5, ENSOARG00000001138, PPM1K* |
| 19 | 6 | 69010001 | 70090000 | 1.079 | 4 | *PDGFRA, LNX1, GSX2, CHIC2* |
| 20 | 6 | 116390001 | 116730000 | 0.340 | 3 | *GAK, CPLX1, ENSOARG00000016536* |
| 21 | 6 | 115960001 | 117040000 | 1.079 | 5 | *LETM1, NELFA, NSD2, SLC49A3, PDE6B* |
| 22 | 7 | 55870001 | 56140000 | 0.270 | 4 | *DMXL2, ENSOARG00000020974, GLDN, CYP19* |
| 23 | 10 | 7260001 | 7360000 | 0.100 | 1 | *ENSOARG00000006632* |
| 24 | 10 | 7310001 | 7650000 | 0.340 | 1 | *ENSOARG00000006641* |
| 25 | 11 | 10750001 | 10940000 | 0.190 | 2 | *INTS2, BRIP1* |
| 26 | 11 | 18250001 | 18490000 | 0.240 | 4 | *NF1, EVI2A, EVI2B, OMG* |
| 27 | 11 | 26350001 | 26540000 | 0.190 | 11 | *ALOX12, RNASEK, U6, ENSOARG00000024277, BCL6B, SLC16A13, SLC16A11, CLEC10A, ASGR2, ASGR1, ENSOARG00000022633* |
| 28 | 11 | 27010001 | 27200000 | 0.190 | 11 | *DNAH2, NAA38, KDM6B, TMEM88, CYB5D1, CHD3, ENSOARG00000022732, RNF227, KCNAB3, TRAPPC1, CNTROB* |
| 29 | 11 | 37190001 | 37430000 | 0.240 | 5 | *CALCOCO2, TTLL6, HOXB13, HOXB9, HOXB7* |
| 30 | 13 | 38660001 | 38870000 | 0.210 | 4 | *RIN2, NAA20, CRNKL1, CFAP61* |
| 31 | 13 | 53290001 | 53660000 | 0.370 | 13 | *ZBTB46, ZGPAT, ARFRP1, STMN3, GMEB2, FNDC11, SRMS, PTK6, EEF1A2, KCNQ2, CHRNA4, ARFGAP1, YTHDF1* |
| 32 | 18 | 50910001 | 51100000 | 0.190 | 0 | *-* |
| 33 | 19 | 23180001 | 23290000 | 0.110 | 1 | *CNTN4* |
| 34 | 22 | 27390001 | 27580000 | 0.190 | 0 | *-* |
| **Total** | | | | | **122** |  |
